# Interpretable representation learning for 3D multi-piece intracellular structures using point clouds

**DOI:** 10.1101/2024.07.25.605164

**Authors:** Ritvik Vasan, Alexandra J. Ferrante, Antoine Borensztejn, Christopher L. Frick, Nathalie Gaudreault, Saurabh S. Mogre, Benjamin Morris, Guilherme G. Pires, Susanne M. Rafelski, Julie A. Theriot, Matheus P. Viana

## Abstract

A key challenge in understanding subcellular organization is quantifying interpretable measurements of intracellular structures with complex multi-piece morphologies in an objective, robust and generalizable manner. Here we introduce a morphology-appropriate representation learning framework that uses 3D rotation invariant autoencoders and point clouds. This framework is used to learn representations of complex multi-piece morphologies that are independent of orientation, compact, and easy to interpret. We apply our framework to intracellular structures with punctate morphologies (e.g. DNA replication foci) and polymorphic morphologies (e.g. nucleoli). We systematically compare our framework to image-based autoencoders across several intracellular structure datasets, including a synthetic dataset with pre-defined rules of organization. We explore the trade-offs in the performance of different models by performing multi-metric benchmarking across efficiency, generative capability, and representation expressivity metrics. We find that our framework, which embraces the underlying morphology of multi-piece structures, facilitates the unsupervised discovery of sub-clusters for each structure. We show how our approach can also be applied to phenotypic profiling using a dataset of nucleolar images following drug perturbations. We implement and provide all representation learning models using CytoDL, a python package for flexible and configurable deep learning experiments.

## Introduction

A central goal of cell biology is to understand the spatial and dynamic organization of the components within the cell and how their interactions contribute to cell function. Enabled by advances in imaging methods, we are now at the dawn of the big data era for cellular imaging^1–4^, in which unprecedented amounts of rich image datasets can enable quantitative characterization of cellular organization and its connections with cellular phenotype.

The term cellular organization encompasses multiple aspects of a cell’s configuration that must be unpacked before further discussion. Here we focus on two of these aspects: spatial protein distributions and shape of multi-piece intracellular structures. For example, the spatial pattern of fluorescently labeled PCNA, representing the punctate morphology of DNA replication foci, changes throughout the cell cycle, making it difficult to quantify due to its dynamic and complex nature. These types of spatial distributions are usually analyzed via the texture patterns they represent, for example computing Haralick texture features^5^. However, the biological meaning of some of these features, such as the *second angular moment of texture*, is difficult to understand. Therefore, for spatial protein distributions, we face the challenge of developing a robust and generalizable analysis workflow that facilitates biological interpretation.

On the other hand, major organelles or subcellular structures can often be analyzed by segmentation, which separates the foreground signal from the background. Intracellular structures composed by a single segmented piece, such as the cell itself or the nucleus, can then be studied via a range of features including, among other methods, shape decomposition using spherical harmonic expansion^1,6,7^. This approach is, however, mainly used for cell and nuclear shapes because it is limited to continuous shapes, and does not easily apply to complex, multi-piece structures like the Golgi apparatus, which has a discontinuous shape. In fact, most intracellular structures exhibit a polymorphic morphology consisting of multiple pieces, which presents an additional challenge for interpretable image analysis pipelines. While each individual piece could be segmented and measured, the entirety of the multi-piece structure cannot be easily represented as a whole.

To overcome these two challenges, we demonstrate the use of 3D point clouds to encode biological data in microscopy images, combined with an unsupervised *representation learning* framework for single cell feature extraction. Representation learning is a field of machine learning that has become an increasingly popular way to extract meaningful features directly from raw data without the need for hand-engineered features^8,9^. These features are in the form of latent variables learned by neural networks during training, which we refer to as *representations*.

An important aspect of the proposed learning framework is that it is generative, meaning we can switch between the original point clouds and the single-cell representations learned, resulting in highly interpretable features and addressing the first challenge described. A key contribution of this work is the use of point clouds to incorporate intensity information present in large 3D images representing spatial protein distribution in a segmentation-free manner. Another important contribution is the adaptation of the point cloud-based approach to handle segmented multi-piece shapes. This is achieved using the concept of signed-distance field, allowing us to generalize our framework to more complex intracellular structures, thus overcoming the second challenge described.

The representations learned by neural networks normally depend on the orientation of an object in the image. Despite the fact that the orientation of the cells is important in many contexts, such as when cells are subject to shear stress, during development or direct migration, it may not be of any biological relevance in other contexts. For example, the orientation of a cell in a monolayer colony may merely reflect the orientation of that colony relative to the microscope stage and not anything biological. Therefore, it would be desirable to design analysis workflows where the image orientation can be factored out of the learned representations if appropriate. We achieved this by leveraging the notion of 3D rotation invariance to extract features that do not depend on an object’s orientation. The incorporation of geometric information in the form of the object orientation into the representation learning process is an example of *geometric deep learning*^10^. By using point clouds as an unifying way of encoding image data, we are able to overcome the challenges described above and take advantage of previous implementations for rotation-invariant feature learning^11–13^ while extending their applications to quantitative cell biology.

Here, we first develop a rotation invariant representation learning framework that uses point clouds to encode relevant information about the underlying biological data. We then use a synthetic dataset of punctate structures to confirm that rotation invariant representations are not sensitive to data orientation and are more compact when learned from point clouds compared to images. We show how 3D rotation invariant features learned from point clouds can be used to recover unique morphological changes of DNA replication foci across the cell cycle without supervision. We also explore the localization patterns of multiple punctate structures and discover novel patterns of intracellular organization. Next, by adapting our framework to handle more complex multi-piece structures, we systematically characterize sources of shape variation of other major intracellular structures like nucleoli, Golgi, and lysosomes. Finally, we demonstrate how the learned representations based on this framework can be used for detecting morphological alterations in a nucleolar drug perturbation dataset, and for visualizing the average phenotype for each drug to aid interpretability.

## Results

### A novel morphology-appropriate framework for learning 3D rotation invariant representations of complex intracellular structures and holistic model evaluation

The 3D rotation invariant representation learning framework has two main components. The first consists of encoding the raw single-cell image data into a point cloud. This encoding process is done in a morphology-appropriate manner for punctate structures and polymorphic intracellular structures, such as DNA replication foci and nucleoli, respectively. The biological meaning of shape differs between these two types of morphologies; we focus on representing the relative location of individual pieces in punctate structures (Fig. 1a), while both relative location and shape of individual pieces are considered important for polymorphic structures (Fig. 1b).

**Figure 1.**
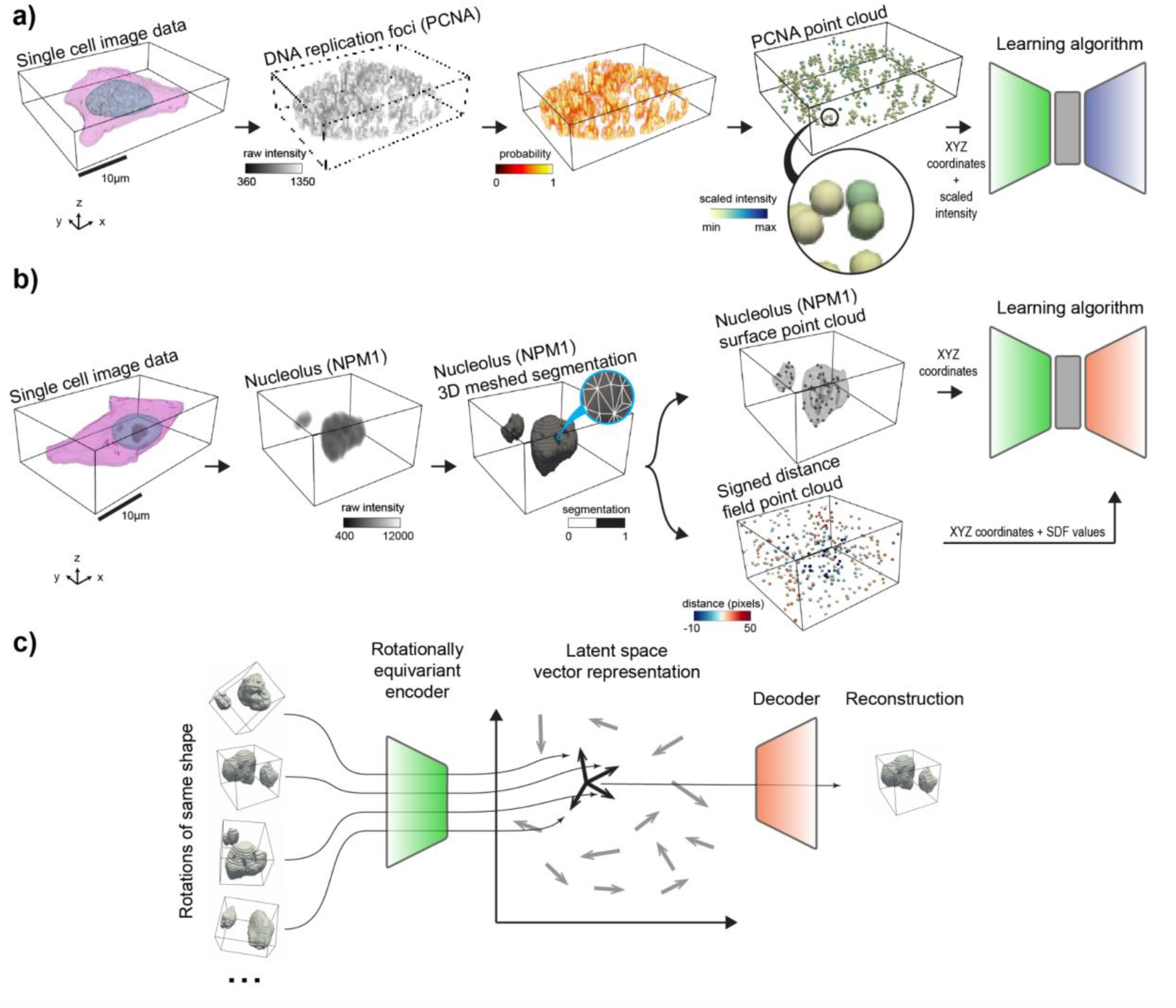
Application-appropriate representation learning framework for complex intracellular structure morphologies. **a)** Example of a punctate structure using DNA replication foci. Shown are single-cell segmentations for the nucleus and cell membrane, and raw intensities for DNA replication foci (via PCNA). 4D (XYZ + intensity) point clouds are sampled from the intensity images by converting intensities to probabilities. The intensity co-ordinate is scaled to ensure that the range of intensity values is similar to the range of XYZ coordinate values. This 4D point cloud is then used as input to the representation learning algorithm. **b)** Example of a polymorphic structure using granular component (GC) nucleoli (via nucleophosmin). Shown are single-cell segmentations for nucleus, cell membrane, and nucleoli (GC). Nucleoli segmentation from single-cell data is used to generate a 3D mesh. A surface point cloud is sampled from the nucleolar mesh. Another point cloud is sampled from the 3D bounding box volume and its points are assigned local signed distance field (SDF) values relative to the surface of the nucleolar mesh. These two point clouds are used by the representation learning algorithm. **c)** 3D rotation invariant representation learning framework using autoencoders. Rotations of the same shape are projected into vector representations using a 3D rotation equivariant encoder. This vector representation is then used to reconstruct the original shape. We take the norm of the vector representation to compute a rotation invariant representation. We do this during training for punctate structures, and after training for polymorphic structures (*Section 4.2 of Methods*).

The second part of the framework consists of a neural network model that consumes the generated point clouds for learning 3D rotation invariant representations of the biological data (Fig. 1c). We define rotation invariance using the group of all rotations in 3D. All the neural network models are designed as *autoencoders*^14^: First, an *encoder network* compresses the generated point clouds into vector latent representations. Next, these latent representations are used by a *decoder network* to reconstruct the input data.

To evaluate the utility of the 3D point cloud encoding, we performed benchmarking against traditional methods using neural network models trained on 3D images directly. We trained classical (rotation dependent) and rotation invariant versions of both image- and point cloud-based models to evaluate the impact of adding the geometric constraint of rotation invariance. We expected point cloud-based models to outperform image-based models for two reasons. First, point clouds are a less redundant way of representing sparse multi-piece intracellular structures compared to image-based models. Second, image-based autoencoders often generate blurry reconstructions that can be particularly problematic for small objects^15,16^. More details about all models used in this paper can be found in *Section 4 of Methods*.

We used a multi-metric approach to evaluate our models and representations. Our goal is to increase transparency behind the performance of these models, and to explore trade-offs. Importantly, we hope to identify models that are quantitatively useful across a broad set of tasks to make gaining biological insight from the learned representations more likely, not necessarily the model that is best for any one metric.. The models were evaluated with respect to their *efficiency*, *generative capabilities* and *representation expressivity* as detailed in Figure S1 (also in *Section 6 of Methods*).

This multi-metric evaluation approach tested different models beyond their ability to reconstruct the input, which is the primary task of autoencoders. For example, this included efficiency metrics that gauged the compute resources and time required to evaluate each model, including the model size, inference time and carbon emissions. We also included generative metrics like the evolution energy between two sample shapes (Fig. S1b). In addition, we also evaluated the models’ expressivity by quantifying the biological information content in the learned representations. Specifically, we used representations for classifying or predicting (via regression) biologically-relevant measurements in an application-appropriate manner (*Section 6.2 of Methods*). For example, we used the representations to predict volume or number of pieces of a given intracellular structure when these properties are relevant for the application. Considering all these metrics together, we quantified the holistic utility of each model (Fig. S1c) and the pros and cons of using each approach.

To aid reproducibility and empower researchers with the ability to independently test and apply these models to their own data, we provide all the representation learning models used in this study via CytoDL, a Python package for configurable 2D and 3D image-to-image deep learning transformations and representation learning, available at https://github.com/AllenCellModeling/cyto-dl. CytoDL is designed consistent with FAIR^17^ practices, and is built to work for diverse use cases, thus making it robust, modular, and flexible to the evolving nature of research.

### 3D rotation invariant point cloud models are efficient, produce low rotation invariance errors and generate good reconstructions in a synthetic dataset of punctate structures

We started by evaluating the effectiveness of 3D rotation invariance and the choice of using point clouds to encode punctate structures using synthetic data. We used cellPACK to create a synthetic dataset of punctate structures with known rules of organization. cellPACK generates 3D models of complex biological environments using novel packing algorithms^18^. To create the synthetic dataset, we used multiple spatial rules for packing spheres in real 3D nuclear shapes based on gradient algorithms (*Section 1.3 of Methods*). These algorithms include a rotationally-dependent *planar gradient* where spheres are packed away from a plane oriented at 0-, 45-, or 90-degrees about the z-axis, and a set of three rotationally-independent gradients including a *radial gradient* where spheres are packed with a gradient away from the centroid of the nucleus, a *random gradient* where spheres are packed randomly, and a *surface gradient* where spheres are packed close to the nuclear boundary, resulting in six total packing rules (Fig. 2a). We packed spheres according to each of these six rules into 254 real nuclear shapes selected from the DNA replication foci dataset (*Section 1.1 of Methods*). Importantly, the stochastic nature of the packing algorithm generates heterogeneity in the distribution of spheres across the simulated nuclei, thus making the recovery of rules via unsupervised learning more challenging, and evaluating the robustness of each representation learning method.

**Figure 2.**
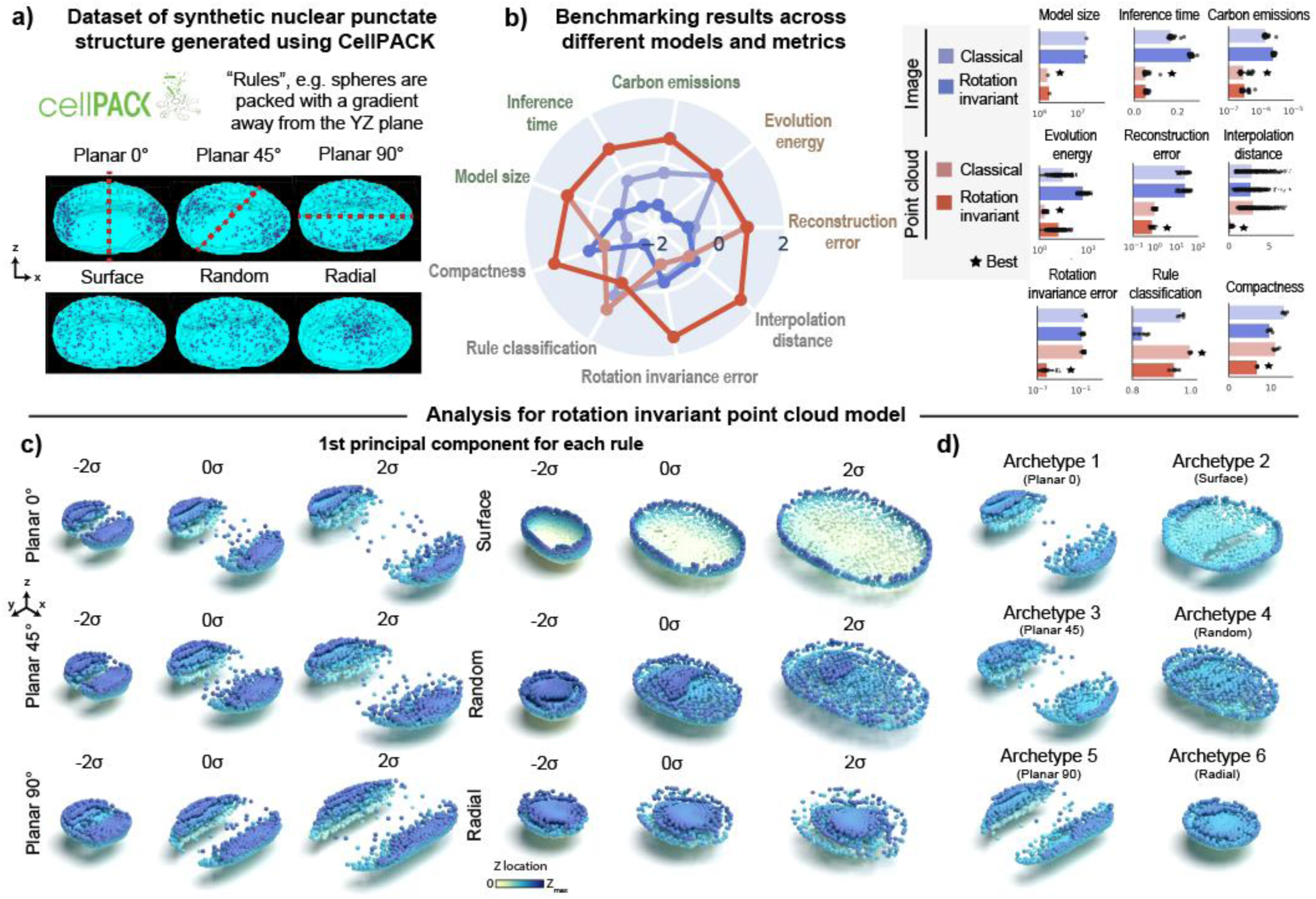
3D rotation invariant point cloud models are efficient, produce low rotation invariance errors and generate good reconstructions in a synthetic dataset of punctate structures. **a)** Dataset of synthetic punctate structures generated using cellPACK. A 3D nuclear shape is packed with 6 different rules. This includes Planar 0, Planar 45, Planar 90, Radial, Random, and Surface. Planar rules pack spheres with a gradient away from a plane indicated in red. Radial rule packs spheres close to the centroid. Random rule packs spheres randomly across the 3D nuclear volume. Surface rule packs spheres close to the nuclear boundary. Each rule is used to pack 254 different nuclear shapes. **b)** Benchmarking unsupervised representations across different models and metrics. (Left) Polar plot showing performance of classical and rotation invariant image and point cloud models across efficiency metrics (model size, inference time, emissions), generative metrics (reconstruction, evolution energy), and representation expressivity metrics (compactness, classification of rules, rotation invariance error, average interpolate distance). Metrics are z-scored and scaled such that larger is better (Right) Bar plots showing raw metric values across models for each metric. Error bars are standard deviations. Best model for each metric is indicated. **c)** 1st principal component for each rule using the rotation invariant point cloud model trained with jitter augmentations. PCA is fit to representations of each rule separately. Shown are normalized PCs (standard deviation (s.d.), σ, units) sampled at 3 map points (-2σ to 2σ in steps of σ). **d)** 6 archetypes computed from the rotation invariant point cloud representations. Each archetype corresponds to one of the 6 rules. All reconstructions shown are cut at mid-plane. Color associated with each point is the distance from the mid-plane in Z.

Since 3D rotation is an important variable associated with the planar rules, we expected 3D rotation invariant models to give us the most compact representations by factoring out this variable. We additionally hypothesized that point cloud models (Fig. 1a) would provide better representations than image models since they better describe the punctate nature of the synthetic data represented by the centroid of the packed spheres (Fig. 2a). To test this hypothesis, we trained two classical and two 3D rotation invariant models using images and point clouds as input data respectively (*Sections 2.1, 3.1, and 4 of Methods*).

We found that point clouds were more efficient across all efficiency metrics (Model size, Inference time and Emissions in Fig. 2b). In addition, point clouds also produced better reconstructions (“Reconstruction error”) and had low evolution energy scores, meaning that the interpolations between two shapes are smooth. Next, we tested whether the implementations of the rotation invariant models were indeed generating representations that were not sensitive to orientation of the input data. We confirmed that this was the case as shown by similar orientation of reconstructions in the last row of Figure S2. We found that rotation invariant representations from point clouds were more compact using the Levina-Bickel intrinsic dimensionality metric (“Compactness”)^19^ and had much lower rotation invariance errors compared to its image-based counterpart (Fig. 2b, *Section 6 of Methods*). All four models were able to reconstruct the unique morphologies associated with each packing rule (Fig. S3). However, we found that representations from both rotation invariant models were slightly worse than their classical counterparts at classifying the six rules (“Rule classification” in Fig. 2b). This was an expected outcome since rotation is an important distinguishing feature of the planar rules and rotation invariant representations are insensitive to this feature. Overall, the 3D rotation invariant point cloud model was an efficient generative model that learned compact and orientation-independent representations for synthetic punctate structures.

Having established the holistic utility of the rotation invariant point cloud model on synthetic data, we next performed principal component analysis (PCA; S*ection 7.1 of Methods*) on the learned representations using this model to interpret their meaning. We performed this PCA on a version of this model trained with jitter augmentations to improve reconstruction quality (Fig S4; also see jitter details in *Section 5.2 of Methods*). By visualizing the first principal component (PC1) of the reconstructions for each rule via a latent walk, we found that PC1 recovers how nuclear size affects each rule’s packing (Fig. 2c). Notably, the rotation invariant reconstructions for all planar rules are aligned in the same direction, allowing us to focus on the subtle differences in spatial distribution between 0- and 45-degrees orientation.

Next, we performed an archetype analysis^20^ to find extremal points in the representations of the synthetic dataset (*Section 7.2 of Methods*). Archetypes are determined so that observations can be approximated by convex combinations of the archetypes. By setting the number of archetypes to six, we found each archetype represented one of the six rules used in cellPACK to generate the synthetic dataset. These results show that the obtained point cloud rotation invariant representations can enable unsupervised rule discovery for a synthetic dataset of punctate structures.

### Rotation invariant point cloud representations recover cell cycle-dependent spatial patterns of DNA replication foci

After establishing its applicability to synthetic data, we tested the representation learning framework on a real single-cell image dataset of punctate structures for biological discovery and hypothesis generation. The dataset contains single-cell images of DNA replication foci in hIPS cells expressing fluorescently tagged PCNA (N=2,420, *Section 1.1 of Methods*). DNA replication foci are punctate and display a continuous change in their overall localization pattern and intensity throughout cell cycle^21^ (Fig. 3a). Due to tagged PCNA fluorescence intensity being an important source of variation for DNA replication foci patterns, we adapted the point cloud sampling strategy so that the raw image intensity is treated as a fourth coordinate, in addition to the XYZ spatial coordinates (Fig. S5a-b, and *Section 3.1.2 of Methods)*. This additional coordinate ensures that intensity information is captured in the learned representations without impacting the rotation invariance of the XYZ spatial coordinates (Section 4.2 of Methods).

**Figure 3.**
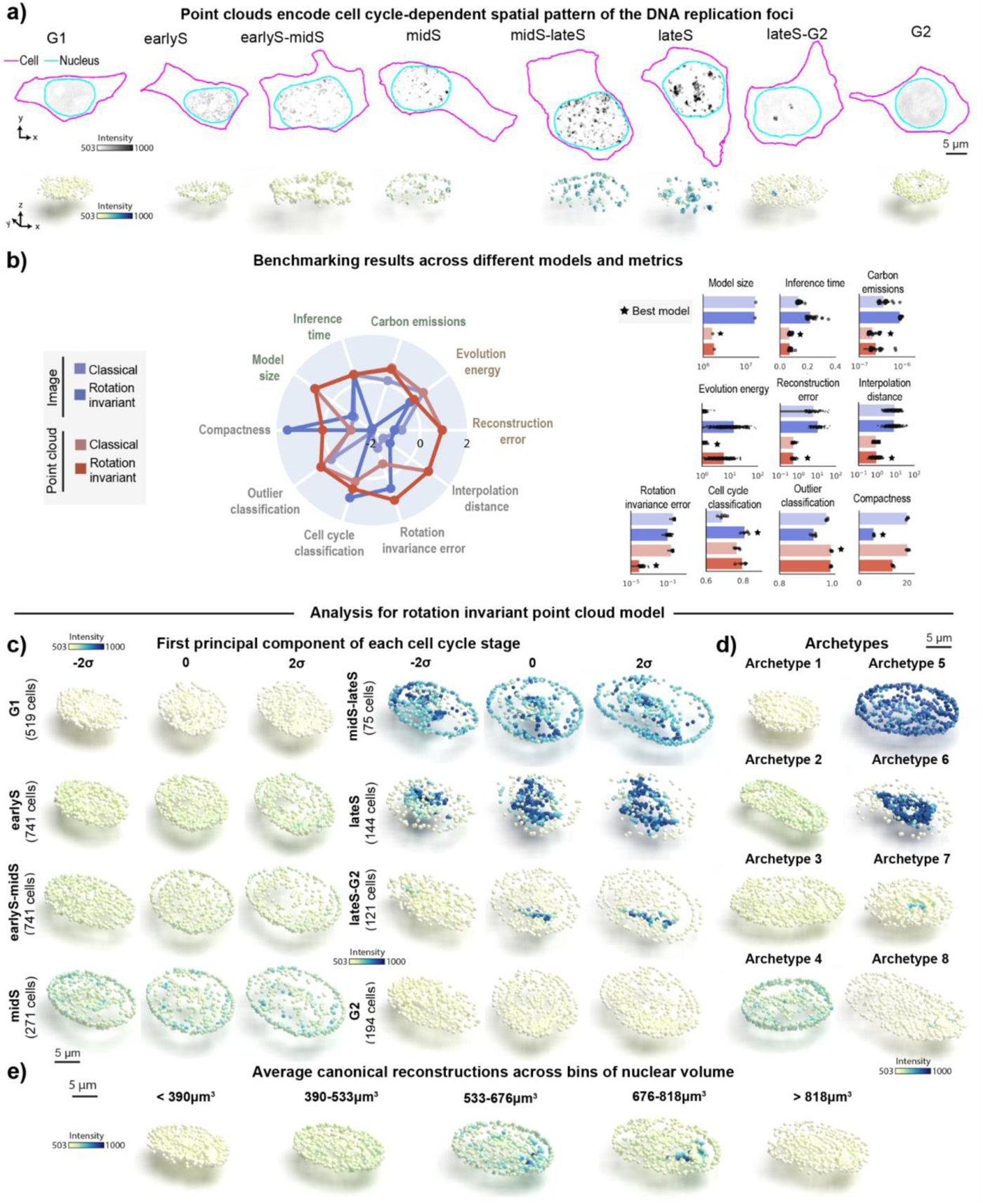
Rotation invariant point cloud representations recover cell cycle-dependent spatial pattern of DNA replication foci. **a)** Dataset of DNA replication foci in hIPS cells expressing mEGFP-tagged PCNA. DNA replication foci have a stereotypical cell cycle dependent localization pattern. Shown are examples of image and sampled point cloud center slices with adjusted contrast for eight expert-annotated cell cycle stages. The scale bar is 5 μm. **b)** Benchmarking unsupervised representations across different models and metrics. (Left) Polar plot showing performance of classical and rotation invariant image and point cloud models across efficiency metrics (model size, inference time, emissions), generative metrics (reconstruction, evolution energy), representation expressivity metrics (compactness, classification of cell cycle via top-2 classification accuracy, rotation invariance error, average interpolate distance). Metrics are z-scored and scaled such that larger is better (Right) Bar plots showing raw metric values across models for each metric. Error bars are standard deviations. Best model for each metric is indicated. **c)** 8 archetypes identified using rotation invariant point cloud representations. Each archetype corresponds to one of the 8 expert-annotated cell cycle stages. **d)** 1st principal component for each cell cycle stage using rotation invariant point cloud model. PCA is fit to representations of each cell cycle stage separately. Shown are normalized PCs (standard deviation (s.d.), σ, units) sampled at 3 map points (-2σ to 2σ in steps of σ). **e)** Average canonical reconstructions across five bins of nuclear volume (*Section 7.3 of Methods*). All reconstructions shown are center slices.

To test whether the representations learned with this data capture biologically relevant features about DNA replication foci localization, we manually classified each single-cell image in this dataset into one of eight cell-cycle stages based on the spatial pattern of PCNA (*Section 1.1 of Methods*). We also manually labeled cells as outliers if they were dead, dying, or did not express PCNA. Next, we used the representations learned by each of the four models to benchmark their performance on various tasks, including classification of cell cycle stages and detection of outliers from the DNA replication foci dataset.

We found that point cloud models were more efficient but not as compact as the rotation invariant image model (Fig. 3b). Point cloud models also provided better overall reconstructions compared to image models (compare reconstructions in Fig. S6f-h to Fig. S6b-c). Despite the poor reconstruction of both image models (Fig. S6b-c), we found that the rotation invariant image model was the best at classifying cell cycle stages (“Cell cycle classification” in Fig. 3b, 81% accuracy vs 80% accuracy for the best point cloud model). This result in particular demonstrates the limits of evaluating models using a single metric alone. We confirmed that poor reconstructions of image models were not due to dataset size or image normalization issues using an alternative approach (Fig. S6d and *Section 4.1.1 of Methods*). We also found that point cloud models had slightly better performance detecting outliers compared to image-based models (“Outlier classification” in Fig. 3b, ∼100% accuracy vs 98% accuracy for the best image model). Finally, we found that the rotation invariant point cloud model had lower rotation invariance error scores compared to its image counterpart. Overall, the results elucidate the challenge of reconstructing sparse intracellular structures using classical image autoencoders^24^, and highlight the good performance of the rotation invariant point cloud representations across many metrics evaluated for the DNA replication foci dataset.

To interpret the representations learned by the rotation invariant point cloud model per cell cycle stage, we performed PCA on this dataset stratified by the manual annotations of cell cycle stages. A latent walk along PC1 for each cell cycle stage revealed some overlap in the morphology and intensity of DNA replication foci between neighboring cell cycle stages (earlyS σ=2 and earlyS-midS σ=-2, for example). This highlights the inherent uncertainty that is present in the task of manual annotation of a continuous process in discrete classes.

We found that an archetype analysis with eight archetypes was able to recover expected cell cycle patterns of DNA replication foci (order of archetypes in Fig. 3d resembles examples in Fig. 3c for σ=0). The archetypes capture three main sources of variation in the dataset as expected. First, overall nuclear shape which is mainly represented by archetypes displaying different elongations. In addition to nuclear shape, the intensity and localization of DNA replication foci are significantly different between archetypes. These two sources of variation seemed correlated as we observe in real PCNA images. Lastly, the spatial pattern of PCNA changes from a dim signal uniformly distributed in the nucleus at G1 to compact bright spots well localized in lateS.

Next, we asked if the learned representations could be used for interpreting the evolution of the DNA replication foci spatial pattern as a function of the cell cycle stage. To test this, we binned nuclear volume into five bins to create a pseudo cell cycle axis (*Section 7.3 of Methods*), and we averaged the rotation invariant point cloud representations of all cells within each bin (Fig. 3e). We observed a continuous orientation-invariant transition that recapitulates the coalescence of uniformly distributed dim DNA replication foci puncta in G1 into a few bright dots in late S. This pattern is followed by signal sparsification into uniform dim punctate again in G2. We observed a moderate distinction between spatial patterns in Figure 3e, which suggests that nuclear size is only a weak proxy for cell cycle stages. These results demonstrate that the learned point cloud rotation invariant representations can recover the overall behavior of DNA replication foci in an unsupervised manner.

### Representation learning framework reveals interpretable spatial patterns for other punctate structures from the WTC-11 hiPSC Single-Cell Image Dataset v1

To assess whether our approach would generalize to other intracellular structures with punctate morphology, we analyzed a larger dataset of punctate structures from the WTC-11 hiPSC Single-Cell Image Dataset v1 (*Section 1.2 of Methods*). This dataset comprises centrioles (N=7.575), peroxisomes (N=1,997), endosomes (N=2,601), nuclear pores (N=17,703), nuclear speckles (N=2,980), cohesins (N=2,380), and histones (N=15,875). Examples of these structures are shown in Figure 4a. Once again, we trained classical and rotation invariant image-and point cloud-based models on this larger dataset (Fig S5c, *Sections 2.1 and 3.1 of Methods)*. In addition to the usual set of evaluation metrics, we tested the applicability of the learned representations for two classification tasks. The first task focused on identifying the specific intracellular structure from the seven options available in the dataset. The second task involved classifying cell-cycle stages (interphase or mitosis) based on the annotations provided within the dataset (*Section 6.2 of Methods*).

**Figure 4.**
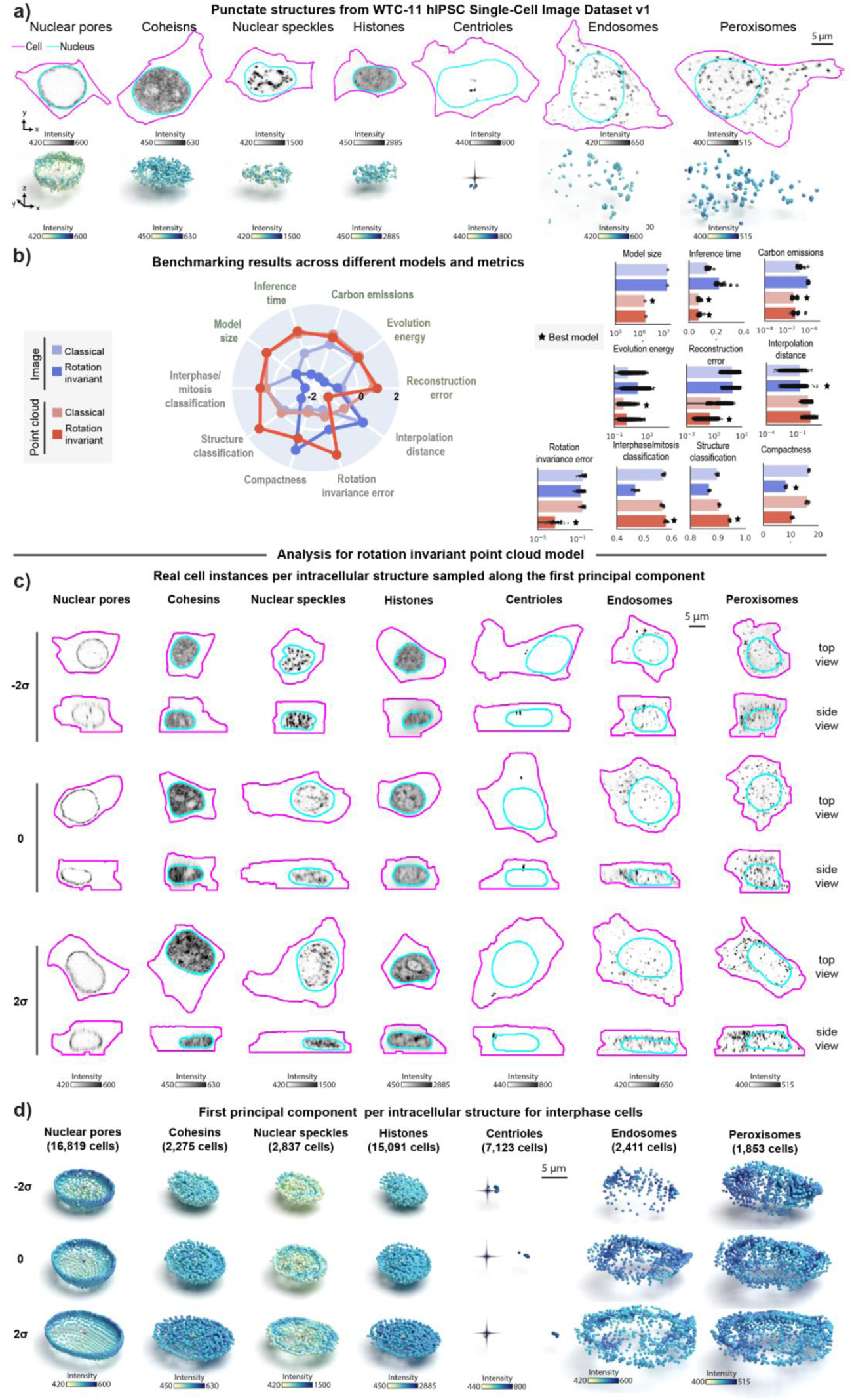
Representation learning framework reveals interpretable spatial patterns for other punctate structures from the WTC-11 hiPSC Single-Cell Image Dataset v1. **a)** Dataset of punctate structures in hiPS cells from the WTC-11 hiPSC Single-Cell Image Dataset v1 including nuclear pores, nuclear speckles, cohesins, histones, centrioles, peroxisomes, and endosomes (Viana 2023). Show are examples of images and sampled point cloud center-slices of the FP-tagged protein. The scale bar is 5 μm. **b)** Benchmarking unsupervised representations across classical and rotation invariant image and point cloud models across efficiency metrics (model size, inference time, emissions), generative metrics (reconstruction, evolution energy), and representation expressivity metrics (compactness, classification, rotation invariance error, average interpolate distance). Classification tasks included classifying 7 different structures, and 6 different interphase/mitotic stages (*Section 6.2 of Methods*). (Left) Polar plot showing performance across models where metrics are z-scored and scaled such that larger is better (Right) Bar plots showing raw metric values across models for each metric. Error bars are standard deviations. Best model for each metric is indicated. **c)** Real examples per map point of PC1 computed using PCA fit to representations of each structure separately using the rotation invariant point cloud model. Only cells in interphase were included. Shown are XY and XZ views. The structure channel is shown as center slices across the nuclear centroid for nuclear pores, cohesins and histones, or as max projections for nuclear speckles, centrioles, endosomes, and peroxisomes. **d)** Latent walk for PC1. Shown are normalized PCs (standard deviation (s.d.), σ, units) sampled at 3 map points (-2σ to 2σ in steps of σ). Reconstructions shown are cut at mid-plane. Membrane centroids are marked for centrioles. Only cells in interphase were considered for this analysis. Centriole reconstructions were rotated to be aligned to the X axis.

Overall, we found that classical image-based models provide better reconstructions when trained with the combination of these seven different punctate structures relative to what we observed for the same models trained on DNA replication foci dataset alone (Fig. S7a-b). Despite this improvement in reconstruction, we noticed that the classical image-based model poorly reconstructs some of these structures including centrioles, peroxisomes, and endosomes (see blurry reconstructions in Fig. S7b). In addition, we found that imposing rotation invariance further deteriorates reconstruction of image models across all structures (Fig. S7c). Once again, an alternative approach confirmed that poor reconstruction was not due to dataset issues (Fig. S7d and *Section 4.1.1 of Methods*).

Both classical and rotation invariant point cloud models produced more accurate and comparable reconstructions, but with spatial distribution artifacts for structures with fewer training samples, like endosomes and peroxisomes, similar to the classical image model (as shown by arrows in Fig. S7f-g). In addition to providing improved reconstructions compared to image models, we found that the rotation invariant point cloud representations performed well at both structure classification (∼95% accuracy vs 90% accuracy for best image model) and cell stage classification (∼58% accuracy vs 57% accuracy for best image model), while being more compact and orientation independent (Fig. 4b).

We then analyzed rotation invariant point cloud representations by computing the first principal component (PC1) for each structure for cells in interphase. To do this, we sampled real single-cell images along PC1 for each structure as shown in Figure 4c. Top and side views in Figure 4c suggest the major source of variation across all seven intracellular structures in this dataset appears to be aspects of cell and nuclear shape like height and elongation. Next, we noticed that centrioles are localized near the nucleus at one extreme of PC1 and gradually migrate towards the cell membrane at the other extreme (column “Centrioles” in Fig. 4c). Additionally, we observed nuclear speckles to be more uniformly distributed within the nucleus at one extreme of PC1 and more concentrated near the nuclear shell forming a ring-like pattern at the other extreme of PC1. Surprisingly, latent walks along PC1 revealed similar patterns as additional sources of variation for these two structures (Fig. 4d). We found that centrioles polarize by moving away from the cell center (represented by a dark cross in Fig. 4d), and nuclear speckles concentrate in a ring-like pattern (last row of column “Nuclear speckles” in Fig. 4d). Overall, these results highlight the ability of our rotation invariant point cloud representations to capture meaningful and biologically relevant variations in the spatial pattern of multiple intracellular structures.

### Rotation invariant representation learning framework generalizes to polymorphic multi-piece structures

We next asked if we could adapt our approach to learn 3D rotation invariant representations for non-punctate intracellular structures, such as nucleoli and Golgi apparatus. These organelles are polymorphic structures where the shape of individual pieces, in addition to the location of these pieces, may be important for the underlying biological process and therefore should be captured by the learned representations^22,23^. We combined the point cloud approach with a signed distance field (SDF^11,24,25^) computed from segmented images, which is a function that represents the signed distance of a position to the nearest part of a shape, to incorporate the shape information of individual pieces into the representation learning framework (Fig. S8 and *Section 3.2 of Methods*). To make this possible, a few modifications had to be made to the autoencoder architecture. First, while the encoder was left unchanged, its input point clouds are now sampled from segmented images instead of raw data, specifically from the surfaces of each of the multi-piece substructures. Next, a second point cloud is sampled from the 3D segmentation volume, and each point is assigned to its local SDF value. We replaced the point cloud decoder with an implicit decoder, which takes the latent representation and spatial coordinates of each point’s location in the second point cloud as input, to reconstruct SDF values at each spatial location (Fig. 1b). Thus, both the encoder and decoder represent the 3D shape in different formats, allowing the model to capture complex shape information using SDFs, while remaining less sensitive to orientation changes. Additionally, for the reasons described in Section 4.2.2 of Methods, we had to relax the generative requirement of the model to make the generalization to polymorphic structures possible.

We applied the adapted framework to images of the granular component (GC) of nucleoli via fluorescently-tagged nucleophosmin (NPM1, N=11,814, Fig. 5 and *Section 1.2 of Methods*) which are part of the WTC-11 hiPSC Single-Cell Images Dataset v1^1^. Nucleoli are multi-compartment condensates that exhibit a broad distribution in both the number of pieces and size^26^ and exhibit rapid rotation in 3D ^27,28^. Given these properties, we expected that 3D rotation invariant representations learned using an implicit definition of the nucleolar surface via an SDF would be more interpretable than representations learned by classical models directly from segmented images. To evaluate this, we trained two classical image models using segmentations and SDFs, two 3D rotation invariant image models using segmentations and SDFs, and one 3D rotation invariant point cloud model using SDFs (*Sections 2.2, 3.2 of and 4 of Methods*). Examples of inputs and outputs of each of these models can be seen in Figure S9.

**Figure 5.**
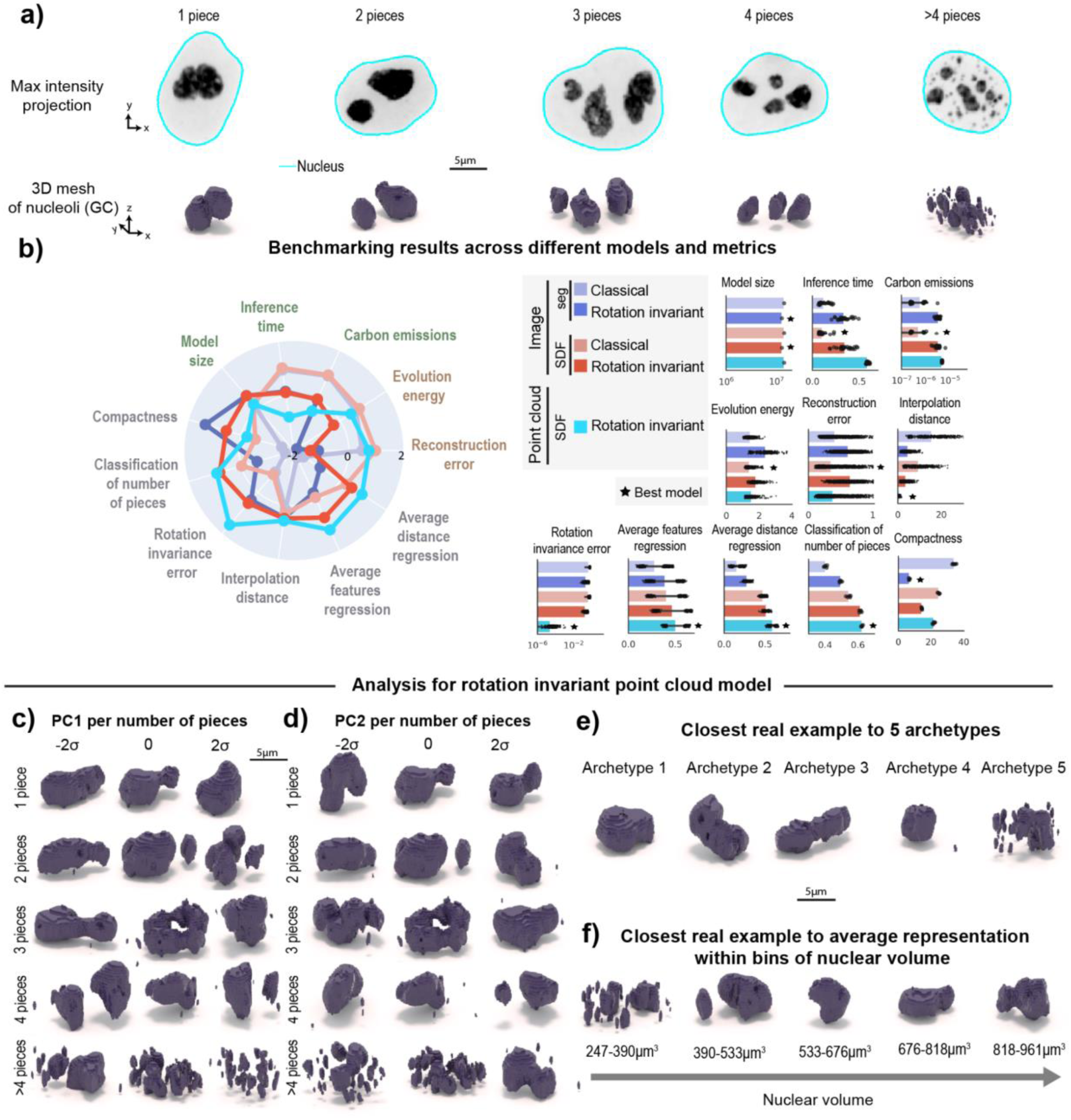
Rotation invariant representation learning framework generalizes to polymorphic multi-piece structures. **a)** Dataset of nucleoli (GC) from the WTC-11 hiPSC Single-Cell Image Dataset v1 (Viana 2023), stratified by number of pieces. Shown are example max intensity projections and corresponding 3D meshes. **b)** Benchmarking unsupervised representations across different models and metrics. (Left) Polar plot showing performance of all models across efficiency metrics (model size, inference time, emissions), generative metrics (reconstruction, evolution energy), representation expressivity metrics (compactness, classification of number of pieces, shape features regression, distance features regression, rotation invariance error, average interpolation distance). Metrics are z-scored and scaled such that larger is better (Right) Bar plots showing raw metric values across models for each metric. Error bars are standard deviations. Best model for each metric is indicated. **c)** 1st principal component for 1 piece, 2 pieces, 3 pieces, 4 pieces, and 5+ pieces examples using rotation invariant point cloud model. PCA is fit to representations of different numbers of pieces separately. Shown are closest real examples to normalized PCs (standard deviation (s.d.), σ, units) sampled at 3 map points (-2σ to 2σ in steps of σ). **d)** 2nd principal component for 1 piece, 2 pieces, 3 pieces, 4 pieces, and 5+ pieces examples using rotation invariant point cloud model. Shown are closest real examples to normalized PCs (standard deviation (s.d.), σ, units) sampled at 3 map points (-2σ to 2σ in steps of σ). **e)** 5 archetypes identified using rotation invariant point cloud model. **f)** Closest real example to average representations of five equal sized bins of nuclear volume (*Section 7.3 of Methods*).

We found that the two classical image models based on segmentations and SDFs, and the 3D rotation invariant point cloud model generate similar quality reconstructions (“Reconstruction error” in Fig. 5b). However, the point cloud model was less efficient in terms of emissions and inference time (Fig. 5b). We also found that 3D rotation invariant image models produce lower quality reconstructions compared to classical image models, as we had observed for models trained on punctate structures (see *Section 6.1 of Methods* for details on how reconstruction error was computed for each model). The results also indicate rotation invariant representations from point clouds are more orientation-independent compared to representations learned from both segmentations and SDFs (“Rotation invariance error” in Fig. 5b). Next, we asked which representations would capture more relevant morphological attributes of nucleoli. To answer this question, we used the learned representation to classify the number of nucleolar pieces in the segmented images and to predict the size, surface area and relative distance between pieces (*Section 6.2 of Methods*). We found that rotation invariant point cloud representations performed best on all of these tasks (“Classification of number of pieces”, “Average feature regression” and “Average distance regression” in Fig. 5b), suggesting that these representations contain relevant biological information.

Next, we used PCA on data grouped by number of pieces to interpret the rotation invariant point cloud representations. Since we had to relax the generative capabilities of this model to achieve rotation invariance, we retrieved the closest real cells while performing a latent walk of PC1 (Fig. 5c) and PC2 (Fig. 5d). We found height and elongation to be the major source of variation for single piece-nucleoli (∼30% of the examples in the dataset; N=3,499, explained variance of PC1was 16% and PC2 was 7%). This was confirmed by computing Pearson correlation with structure elongation (r=0.56 for PC1, r=0.05 for PC2) and height (r=0.13 for PC1, r=0.01 for PC2). In the remaining 70% of the dataset (N=8,315), where nucleoli consist of multiple pieces, the predominant source of variation appears to be the distance between pieces and the relative size of these pieces. For example, when considering nucleoli composed of two pieces, we observe PC1 (explained variance was 19%) to represent the height of the larger piece and the size of the small piece (Fig. 5c). In addition, we found both PC1 and PC2 to correlate with the average distance between pieces (r=0.42 for PC1, r=0.17 for PC2). By performing an archetype analysis with five archetypes, we found that three archetypes represent nucleoli with a single piece but different elongations (archetypes 1-3 in Fig 5e). Archetype four represents nucleoli with one large piece and one small piece, which is a common configuration in the dataset, and archetype five represents nucleoli fragmented in many small pieces.

Motivated by previous observations of cell cycle-dependent nucleolar morphology^29^, we asked whether rotation invariant representations would capture nucleolar changes as a function of the cell cycle. We again used nuclear volume bins to create a pseudo cell cycle axis and visualized the closest real example to the average representation within each bin (*Section 7.3 of Methods*). Consistent with previous observations^29^, we found that cells exiting division (small nuclear volume) have nucleoli that are fragmented into multiple pieces that coalesce into a single large piece as the cells grow and progress towards mitosis (Fig. 5f). Altogether, the results show that this representation learning framework can be successfully adapted using SDFs to polymorphic structures and that it provides representations that capture relevant aspects of the nucleolar biology.

### Learned representations enable shape variation profiling on multiple polymorphic structures

We next expanded this method to profile multi-piece shape variation on the granular component (GC) of nucleoli (N=11,814) and three other polymorphic structures. To do this, we analyzed a subset of polymorphic structures from the WTC-11 hiPSC Single-Cell Image Dataset v1 (*Section 1.2 of Methods*). This subset comprised the dense fibrillar component (DFC) of nucleoli (N=9,923), lysosomes (N=10,114), and Golgi (N=6,175). Representative images of each structure are shown in Fig. 6a. We applied the SDF point cloud rotation invariant representation learning framework to this dataset of four different polymorphic structures, and once again benchmarked it against classical and rotation invariant segmentation-based and SDF-based image models (Fig. S8). We focused here on learning scale invariant representations by scaling all intracellular structures on a cell by cell basis (*Section 2.2.2 of Methods*).

**Figure 6.**
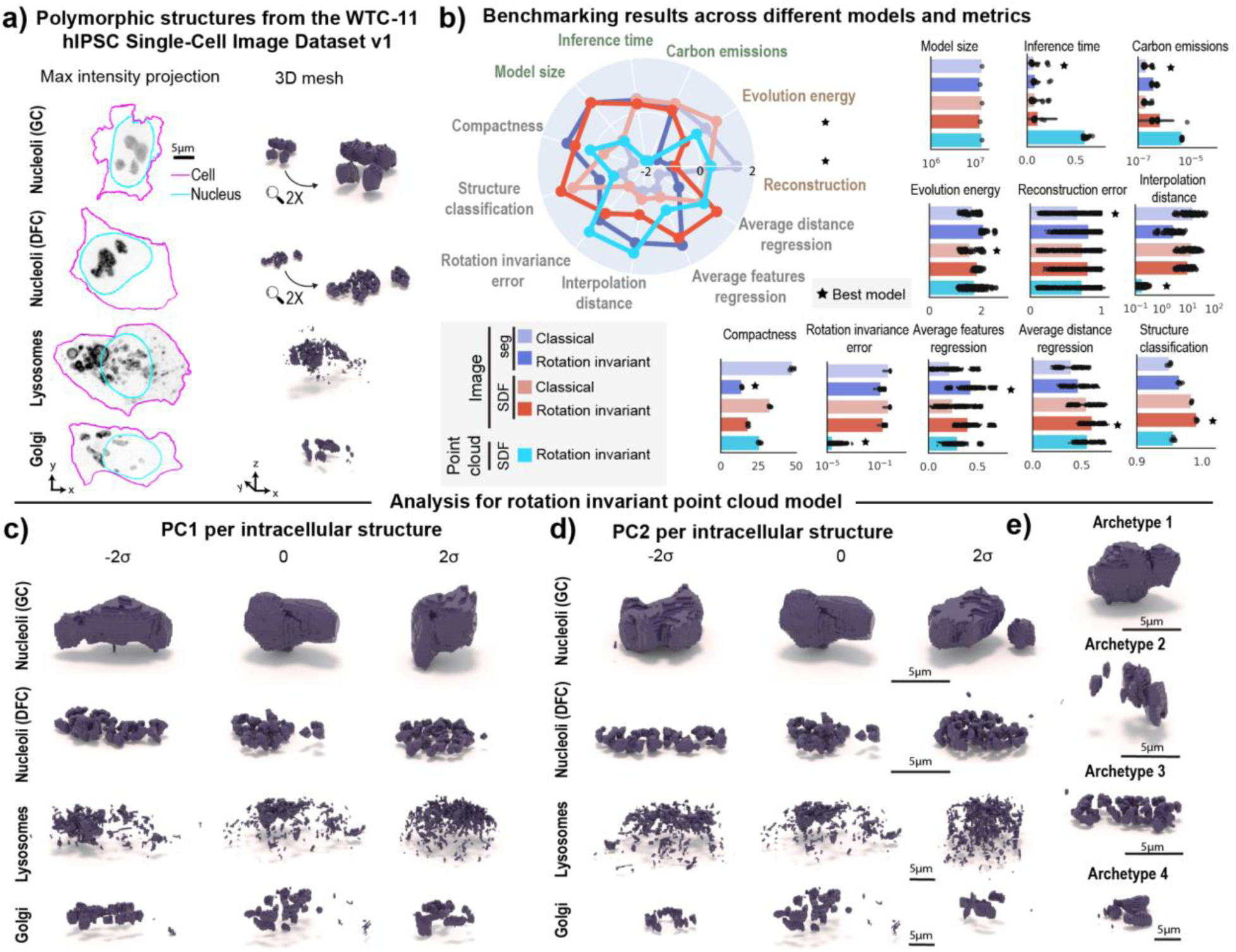
Learned representations enable shape variation profiling on multiple polymorphic structures. **a)** Dataset of nucleoli GC, nucleoli DFC, lysosomes, and Golgi from the WTC-11 hiPSC Single-Cell Image Dataset v1 (Viana 2023). Shown are example max intensity projections and corresponding 3D meshes. **b)** Benchmarking unsupervised representations across different models and metrics. (Left) Polar plot showing performance for all models across efficiency metrics (model size, inference time, emissions), generative metrics (reconstruction, evolution energy), representation expressivity metrics (compactness, classification of number of pieces, shape features regression, distance features regression, rotation invariance error, average interpolation distance). Metrics are z-scored and scaled such that larger is better (Right) Bar plots showing raw metric values across models for each metric. Error bars are standard deviations. Best model for each metric is indicated. **c)** 1st principal component for each structure using rotation invariant point cloud model. PCA is fit to representations of each structure separately. Shown are closest real examples to normalized PCs (standard deviation (s.d.), σ, units) sampled at 3 map points (-2σ to 2σ in steps of σ). **d)** 2nd principal component for each structure using rotation invariant point cloud model. Shown are closest real examples to normalized PCs (standard deviation (s.d.), σ, units) sampled at 3 map points (-2σ to 2σ in steps of σ). **e)** 4 archetypes identified using rotation invariant point cloud representations.

While this scaling strategy prevents us from comparing sizes across different intracellular structures, it helps preserve the resolution of structures occupying only a few voxels. Examples of inputs and outputs of each of the models used can be seen in Figure S10.

We again observed that the rotation invariant point cloud representations continued to display the lowest rotation invariance errors of any model tested (Fig. 6b). We found that all rotation invariant models were more compact than their classical counterparts. We also found that both rotation invariant image models were worse at reconstruction and evolution energy than their classical counterparts (Fig. 6b). Interestingly, all models were able to classify structure equally well (“Structure classification” in Fig. 6b). This suggests that the morphology of these structures is distinct enough to be easily discerned using unsupervised learned representations, despite all being referred to as “polymorphic”.

We performed PCA on rotation invariant representations learned for each of the four intracellular structures and visualized the first two principal components (Fig. 6d-e). We identified features like overall elongation and number of pieces as primary sources of variation for both components of nucleoli, which are known to be nested via liquid-liquid phase separation^30^. In the case of lysosomes, we observed PC1 and PC2 to be related to aspects of cell shape. This was further confirmed by computing Pearson correlation between PC1 (explained variance ∼15%) and cell elongation (r=-0.42), and PC2 (explained variance ∼8%) and cell height (r=0.28). PC1 also displayed strong correlation with the average distance between lysosome pieces (r=-0.45 for PC1), suggesting that the first principal component also captures information about lysosome spatial clustering. In the case of Golgi, the first two principal components (explained variance ∼13% for PC1 and 9% for PC2) appeared to capture aspects of elongation and fragmentation. This was corroborated by computing correlations with structure elongation (r=-0.61 for PC1 and r=0.06 for PC2), the average distance between Golgi pieces (r=-0.55 for PC1 and r=0.27 for PC2), and number of pieces (r=0.14 for PC1 and r=0.05 for PC2). An archetype analysis with four archetypes retrieved two granular component (GC) nucleolar morphologies that represent aspects of elongation (Archetype 1) and number of pieces (Archetype 2). The other two archetypes were examples of the dense fibrillar component (DFC; Archetype 3) of nucleoli and the Golgi (Archetype 4).

Overall, representations from all SDF-based models were able to regress information about distances between pieces better than segmentation-based models, whereas the rotation invariant representations were able to regress information about volume and area of pieces better than classical models. While the point cloud rotation invariant model performed well in generation and representation expressivity metrics, no single model excelled across all metrics. This observation suggests that application-appropriate model selection is key for achieving optimal results. For example, if the goal of an analysis is to learn representations that are not sensitive to rotations, then the rotation invariant point cloud approach would be most appropriate, but if the goal is to learn a model that reconstructs the data well, then a classical segmentation model would be most appropriate in this case.

### Evaluating drug effects on nucleolar morphology using unsupervised representation learning

We then proceeded to test the applicability of the representation learning approach to a perturbation detection task using a drug screening dataset. We imaged WTC-11 hiPS cells expressing an endogenously, fluorescently tagged nucleophosmin, representing the granular component of nucleoli. Cells were treated with 16 different drugs at relatively low concentration to induce subtle phenotypic alterations (*Section 1.4 of Methods*). Analysis was conducted on cells imaged two hours after treatment.

We used the representation learning framework to extract unsupervised representations for cells in the dataset (N=1,025). To do so, we fine-tuned the models trained on the dataset of nucleolar (GC) single-cell images described in the section “Rotation invariant representation learning framework generalizes to polymorphic multi-piece structures”. We followed the methods described in Chandrasekaran et al.^2^ to evaluate the performance of these fine-tuned models. To do so, we computed the *mean average precision* to measure how distinguishable different single cells of a drug-treated set are from untreated cells (DMSO; N=140), and a q-value statistic based on permutation testing.

The results are summarized in Figure 7a, where we plot the q-value per drug for each model. Drugs with q-value under the significance threshold of 0.05 are considered by that model as causing significant alterations in nucleolar morphology. Aside from the first two drugs, we found a significant difference in the behavior of segmentation- and SDF-based models. Therefore, we sorted the x-axis from low to high average q-value based on all SDF models. Consequently, drugs on the left side of the plot induce a stronger phenotypic change compared to drugs on the right side. Figure 7b shows a table with more details about each drug, such as name, concentration, molecular target or mechanism of action, effect based on literature review and effect observed on nucleoli based on visual inspection of this drug dataset. Representative examples of the range of phenotype of each drug are shown in Figure 7c.

**Figure 7.**
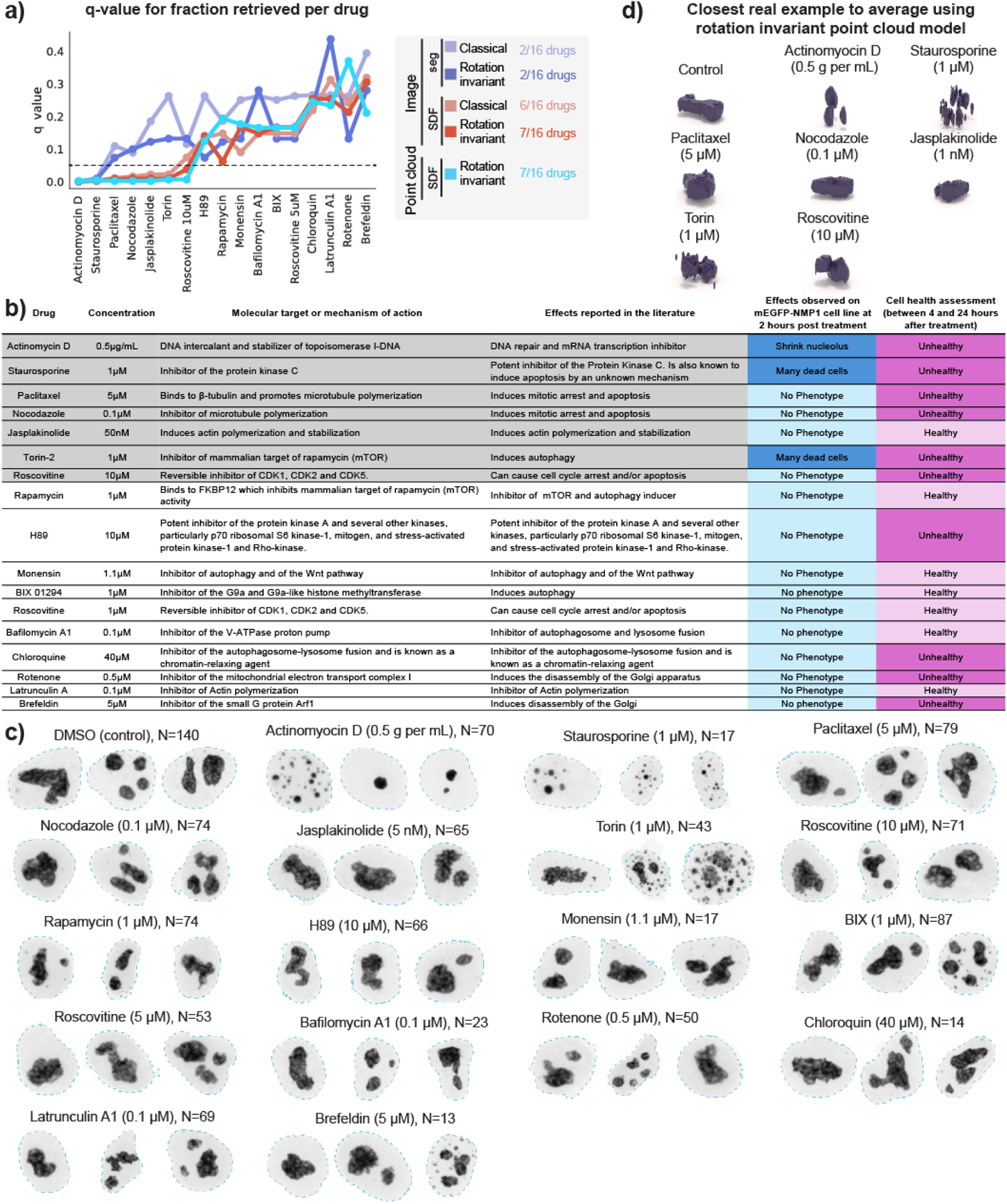
Learned representations allow for morphological profiling of nucleoli under different perturbations. **a)** Q-value statistics^2^ per drug (*Section 1.4 of Methods*) and per model indicating the confidence of each model distinguishing a given drug from control. **b)** Table listing molecular target or mechanism of action, effects reported in the literature, effects observed on the mEGFP-NPM1 cell line at 2 hours post treatment, and cell health assessment between 4 and 24 hours after treatment for each drug. **c)** Three representative examples of nucleoli (GC) for the control (DMSO) and each of the 16 drugs used in this study. **d)** Closest real sample to the average representation for the seven drugs that fall below the q-threshold using the rotation invariant point cloud model.

The first drug to appear on the x-axis of Figure 7a is Actinomycin D, indicating that this drug is the one with the strongest effect on nucleolar morphology. This drug works as a control in this analysis since it is the only well characterized drug in this study that is known to target the DNA and cause significant alteration in nucleolar size, as described in Figure 7b and shown in Figure 7c. Next, we found Staurosporine to have the second strongest effect. A visual inspection of images of cells treated with this drug reveal the presence of many dead cells where nucleoli display very abnormal morphology (Fig 7c). Both Actinomycin D and Staurasporineno were identified as being distinguishable from the control (below q=0.05 threshold) by all models.

No other drug was identified by either the classical or rotation invariant image-based segmentation models. On the other hand, the SDF based models identified several other drugs that could be associated with off-target effects. Starting from the left side of the plot in Figure 7a, these models next identified Paclitaxel and Nocodazole, which were associated with cell death 24 hours after treatment, while also locking cells in mitosis (Fig. 7b-c). SDF-based models also detected Jasplakinolide which is known to promote actin polymerization and is associated with higher nuclear volumes^31^. Visual inspection of the image data did not reveal any apparent nucleolar alteration (Fig. 7c) or change in cell health within 24 hours after treatment, suggesting that the effect of this drug is subtle. Next, Torin was detected by all SDF models, which was observed by visual inspection to induce cell death, like Staurosporine. Lastly, Rascovitine at 10µM was detected only by the SDF rotation invariant models. While there was no visible alteration of nucleoli morphology in this dataset at this concentration, Roscovitine at 10µM caused cell death at 4 hours in images of a different dataset of H2B-tagged cells (Histones, data not shown), and is known to cause nucleolar segregation^32,33^ at higher concentrations. These results suggest that our representation learning framework captures subtle concentration-dependent phenotypes that are not visible by eye. Overall, we observed three different categories of drugs that were retrieved using different models; “sledgehammer” phenotypes that were detected by all models, subtle off-target effects that were detected by SDF models, and subtle concentration-dependent phenotypes that were detected only by the rotation invariant SDF models. The remaining nine drugs in the dataset did not induce visible alterations to nucleolar morphology, although four of them induced cell death at later time points, including H89, Chloroquine, Rotenone and Brefeldin.

Visualization of the closest real example to the average representation using the point cloud model emphasized that Actinomycin D, Staurosporine and Torin-treated nucleoli were visually distinct from the control, confirming that nucleolar morphology is strongly affected by these drugs (Fig. 7d). These results illustrate that our 3D representation learning approach can enable perturbation detection and phenotype profiling, and indicate the need for follow up experiments to confirm the impact of some of these drugs, like Jasplakinolide, on nucleolar morphology.

## Discussion

In this paper, we developed a morphology-appropriate 3D rotation invariant representation learning framework for multi-piece intracellular structures using point clouds. We benchmarked this framework against classical and rotation invariant image-based models using a novel multi-metric evaluation criterion that focuses not only on traditional reconstruction quality but also on measurements that can be important for downstream analysis and biological discovery. We found that our morphology-appropriate approach can be more efficient, can produce meaningful reconstructions, and can learn more compact and expressive representations across a range of tasks than classical image and point cloud models. We applied this framework to synthetic and real single-cell image datasets for punctate structures, like DNA replication foci, and polymorphic structures, such as nucleoli. Our results reveal that geometry-aware choices of encodings and neural network architectures can enable unsupervised discovery and interpretation of variability in the morphology of several multi-piece intracellular structures.

We first demonstrated this principle by recovering the rules used to create a synthetic dataset of punctate structures using cellPACK in a fully unsupervised way. We then recovered expected morphological patterns for different intracellular structures using unsupervised methods, such as archetype analysis and PCA, and by identifying biologically interpretable localization patterns for structures, like DNA replication foci and centrioles. Interestingly, the good performance achieved by different models in classifying cell cycle stages using PCNA suggests that manual annotation of this information can be replaced or assisted by unsupervised representation learning. In addition, the learned representations for centriole capture its repositioning from the cell center towards the periphery, a behavior which is known to be mediated by the microtubule network^34,35^. The learned representations also recapitulate a known axis of morphological change of nuclear speckles, which goes from many, small, irregularly shaped speckles to larger, rounder shaped speckles. This is known to occur when transcription is inhibited in cells and is also the primary axis of variability between cell types^36^.

We further showed how the framework can be extended to polymorphic structures and characterized the full complexity of nucleolar shape variation as an example. When applied to a larger dataset of polymorphic structures, the unsupervised representation learning revealed interesting spatial organization aspects of these structures. For example, Golgi fragmentation captured by the first two principal components of rotation invariant representations is often described as an important morphological feature under both normal and physiological conditions, and has connections with different signaling pathways^23^. However, we observed a trade-off when using the rotation invariant point cloud model where the model produced good reconstructions and learned expressive representations, but was inefficient compared to image models, potentially limiting its ability to scale to much larger datasets and its utility to users with limited computational resources.

Finally, we evaluated the utility of our approach on phenotypic profiling of a nucleoli perturbed image dataset and demonstrated the interpretability of the learned representations. In general, we noticed that segmentation-based models were not able to detect drugs like Torin-2 that caused clear alterations on nucleolar morphology as a result of cell death. This result suggests that SDF encodes information relevant for perturbation detection. It is important to note that our assessment of cell health based on the number of dead cells in the FOV does not distinguish between different processes by which cells can die. This could be a possible explanation for why some drugs that induce death were detected as being distinguishable from the control, while others were not. A larger drug screening single-cell image dataset is necessary to comprehensively validate our findings and evaluate the robustness of our approach.

In this work, we focused the experiments on 3D confocal images of intracellular structures in the hiPS cell, but our framework is general and can be applied to other cell types and imaging modalities. For example, single-molecule localization microscopy (SMLM) naturally generates large 3D point cloud data for single molecule interaction at the 10-20 nm resolution range^37^. Similarly, imaging-based spatial transcriptomics methods that measure a small subset of genes using single-molecule fluorescence in-situ hybridization (smFISH) technologies typically generate point locations of RNA localization^38^, and antibody-based immunofluorescence imaging methods generate protein localization maps in 3D systems such as organoids^39,40^. These datasets are often quantified using point cloud clustering analysis and could benefit from unsupervised representation learning frameworks such as the one proposed here.

Many current techniques for analyzing single-molecule localization microscopy operate in two dimensions, forcing researchers to project their 3D data into 2D for analysis purposes^41,42^. This trend is prevalent in cell biology and medicine, where numerous publicly accessible datasets are predominantly in 2D^2,3,43^. The development of 3D analysis frameworks combined with availability of 3D data, can mitigate barriers to conducting comprehensive 3D analyses across diverse imaging modalities. This, in turn, reduces the risk of artifacts and allows analysis to be centered on the native data formats.

Previous studies have introduced unsupervised representation learning approaches for cell images using autoencoders with geometric deep learning^44,45^. Our work complements these approaches in three ways: first, by incorporating the notion of orientation invariance into our intracellular structure morphology-dependent framework for representation learning; second, by providing a systematic multi-task benchmark to evaluate the utility of each model that goes well beyond traditionally assessed reconstruction quality; third, by focusing our analysis on 3D multi-piece intracellular structures with complex morphology and spatial distribution. Our approach was tested across datasets of different sizes commonly obtained in typical single cell imaging studies, ranging from a few hundred to tens of thousands samples, thus providing a guideline for the quality of unsupervised representations in different cases. For instance, we observed poor image-based reconstructions on small datasets like the DNA replication foci dataset. This observation might be closely related to our deliberate choice of using vanilla autoencoders throughout our analysis, and many modifications of autoencoders have been shown to improve different aspects of reconstruction^46,47^, disentanglement^44,48,49^, causal inference^50,51^, and dynamics^52,53^.

Our framework can be further improved in multiple ways. For example, our results indicate cell and nuclear shape are major sources of variation because that information was not factored out of our learning framework and therefore, become confounding variables. While this reflects a true coupling between cell and nuclear shape and structure localization, alternative approaches may offer a way to decouple these confounding variables from learned representations. For instance, one could incorporate reference information about other intracellular structures for answering questions about intracellular structure co-localization^54^. Another possibility for improving our framework could be adding mutual exclusivity rules to further constrain the models and move towards a better understanding of compartmentalization^15^. Finally, given the limited ability to simultaneously tag and visualize multiple structures^55,56^, extending our framework to predict spatial patterns of a set of structures given the representation from another set could help build a holistic description of intracellular organization. The morphology-appropriate representation learning using point clouds and SDFs that we have described here is flexible and can be modified to incorporate several such improvements.

In summary, we have begun to develop a computational analysis pipeline for interpretable representation learning of complex multi-piece intracellular structures. An important goal of this work is to make the data, models and analysis tools freely available to the community, so that it can serve as a benchmark for further methods development for 3D analysis. We hope that this work can spur the interest of the cell biology community into new ways of analyzing and interpreting complex intracellular organization.

## Acknowledgments

We thank Aditya Nath, Ben Gregor, Calysta Yang, Derek Thirstrup, Joyce Tang, Mackenzie Coston, Melissa Hendershott and Winnie Leung for collecting the perturbation dataset used in this paper. We thank Megan Riel-Mehan, Ruge Li and Graham Johnson for developing cellPACK. We thank Clare Gamlin, Erin Angelini and Serge E. Parent for editing suggestions to the text. The WTC line that we used to create our gene-edited cell lines was provided by the Bruce R. Conklin Laboratory at the Gladstone Institute and UCSF. S.M.R. and C.L.F. were supported for some of this work by the National Human Genome Research Institute of the National Institutes under Award Number UM1HG011593. The content is solely the responsibility of the authors and does not necessarily represent the official views of the National Institutes of Health. This article is subject to HHMI’s Open Access to Publications policy. HHMI laboratory heads have previously granted a nonexclusive CC BY 4.0 license to the public and a sublicensable license to HHMI in their research articles. Pursuant to those licenses, the author-accepted manuscript of this article can be made freely available under a CC BY 4.0 license immediately upon publication. We wish to thank Allen Institute founders, Jody Allen & Paul G. Allen, for their vision, encouragement, and support.

## Author contributions

Conceptualization, R.V., A.F., G.G.P., S.M.R., J.A.T., M.V.P.; Data Curation, R.V., A.F., A.B., C.L.F., N.G., S.S.M.; Formal Analysis, R.V., A.F.; Investigation, R.V., A.F., A.B., N.G., S.S.M., S.M.R., J.A.T., M.V.P.; Methodology, R.V., A.F., J.A.T.; Project Administration, R.V., M.V.P.; Resources, R.V., A.F., B.M.; Software, R.V., A.F., B.M.; Supervision, S.M.R., J.A.T., M.V.P.; Validation, R.V., A.F., A.B.; Visualization, R.V., A.F.; Writing of the original draft, R.V., M.V.P.; Writing review and editing, R.V., A.F., A.B., G.G.P., S.M.R., J.A.T., M.V.P.

## Declaration of interests

The authors declare no competing interests.

## Methods

### 1. single-cell Image Datasets

#### 1.1 DNA replication foci dataset

Spinning-disk confocal 3D images taken of a fluorescently tagged cell line that targets proliferating cell nuclear antigen (PCNA) labeling DNA replication foci with mEGFP were processed to create the DNA replication foci dataset^57^. Fluorescent cell-membrane and DNA dyes tagged the cell boundary and nucleus, respectively. Nuclear segmentations were obtained using the protocol described in Viana et al. ^1^, with the only difference being that nucBlue dye was replaced with nucViolet dye. Segmentations of DNA replication foci were generated for each FOV, using three different segmentation workflows created using the *Allen Cell & Structure Segmenter*^58^ to segment specific DNA replication foci morphologies. Next, we visually identified which segmentation workflow was best for each cell and saved the result in an empty FOV at that cell’s correct location. More details about the dataset and images are available here https://open.quiltdata.com/b/allencell/packages/aics/nuclear_project_dataset_4.

Cells in interphase were labeled by an expert as belonging to one of nine classes - G1, earlyS, early-midS, midS, midS-lateS, lateS, lateS-G2, G2, and unclear. Unclear labels were dropped during analysis. About 3% of cells were labeled as outliers based on bad segmentations of DNA replication foci, cells appearing dead or dying, no EGFP fluorescence, and bad segmentations of cells and nuclei. Dead cells and no fluorescence were used for the outlier detection task, accounting for 16 cells out of a total of 2,420 cells.

#### 1.2 WTC-11 hiPSC Single-Cell Image Dataset v1

Spinning-disk confocal 3D images taken from 25 endogenously tagged hIPS cell lines were processed to create the WTC-11 hiPSC Single-Cell Image Dataset v1^1^. Fluorescent cell-membrane and DNA dyes tagged the cell boundary and nucleus, respectively. Cell, nuclear and structure segmentations were used as provided in the dataset release available here https://open.quiltdata.com/b/allencell/packages/aics/hipsc_single_cell_image_dataset.

We performed analysis on histones via H2B (N=15,875), nuclear pores via Nup153 (N=17,703), peroxisomes via PMP34 (N=1,997), endosomes via Rab-5A (N=2,601), centrioles via centrin-2 (N=7,575), cohesins via SMC1A (N=2,380) and nuclear speckles via SON (N=2,980) as selected punctate structures from this dataset. We selected nucleoli (DFC) via fibrillarin (N=9,923), nucleoli (GC) via nucleophosmin (N=11,814), lysosomes via LAMP-1 (N=10,114), and Golgi via sialyltransferase (N=6,175) as polymorphic structures. While we used all single-cell images for training our models, we limited our analysis to interphase cells.

#### 1.3 cellPACK synthetic single-cell dataset

We used cellPACK to create synthetic point clouds within real nuclear shapes^18^. cellPACK provides an algorithm to create high-resolution 3D representations of the biological mesoscale based on specified rules. Segmentation of 254 randomly chosen nuclei from the DNA replication foci dataset were converted into a triangulated mesh and used as input to cellPACK. cellPACK then packed 256 spheres with a radius of 1 voxel within these meshes based on four distinct rules. **Random**: points were generated uniformly at random inside the nucleus; **Planar gradient rule**: points were generated inside the nucleus with a bias away from a plane. The plane contains the centroid of the nucleus, and its orientation is specified by a normal vector. We used normal vectors with three different orientations: i) θ=0, the normal vector points along the z-axis (0x + 0y + 1z) where the longest axis of the nucleus is the y-axis. ii) θ=45°, the normal vector is (0x + 1/√2y + 1/√2z). iii) θ=90°, the normal vector points along the y-axis; **Surface gradient rule**: points were generated with a strong bias towards the nuclear surface **Radial Gradient**: points were generated with a bias towards the centroid of the nucleus. For each rule, cellPACK generated a point cloud with 256 points for each nucleus shape. This dataset is available for download at https://open.quiltdata.com/b/allencell/tree/aics/morphology_appropriate_representation_learning/cellPACK_single_cell_punctate_structure/

#### 1.4 Drug dataset

A collection of well-characterized drugs was used to perturb the Allen Institute for Cell Science cell line AICS-50 (WTC-11 hiPSC endogenously tagged for mEGFP-NPM1, tagging nucleoli (GC)). Drugs and concentrations were selected because cell treatment with each of them induced a well-characterized effect on one major cellular structure morphology that could be visually observed within 24 hours of treatment (Table 1) and was not associated with massive cell death within the first two hours of treatment, except for Jasplakinolide. Cells were seeded on a 96-well glass bottom plate using the protocol described in Gregor et al.^59^. Four days post seeding 2D brightfield low magnification well overviews were acquired and used for position selection following the same criteria as described in Viana et al.^1^. Following position selection cells were washed once with pre-warm phenol red-free mTeSR then media was replaced with drug-containing phenol red-free mTeSR media at the indicated concentration (Supp table X). The cells were then placed back on the spinning-disk confocal microscope stage where they were maintained at 37°C with 5% CO2 for two hours before the start of imaging at high magnification (120X). Images were acquired with three identical ZEISS spinning-disk confocal microscopes with 10X/0.45 NA Plan-Apochromat (for well overview and position selection) and 100X/0.8 NA Plan-Apochromat (Zeiss) (for high-resolution imaging) and ZEN 2.3 software (blue edition; ZEISS). The spinning-disk confocal microscopes were equipped with a 1.2X tube lens adapter for a final magnification of 12X or 120X, respectively, a CSU-X1 spinning-disk scan head (Yokogawa) and two Orca Flash 4.0 cameras (Hamamatsu). 3D FOV image stack acquisition was performed with two cameras allowing for simultaneous acquisitions of a brightfield and an mEGFP (excited with 4.6mW of a 488nm laser) channel. Exposure time was 100ms. The resulting images were of 16 bits and 924x624 pixels^2^ in xy dimension after 2x2 binning. FOVs had a final xy pixel size of 0.108 μm and z-stacks composed of 100 z-slices (to encompass the full height of the cells within an FOV) acquired at a z interval of 0.29μm. Transmitted light (bright-field) images were acquired using a red LED light source with a narrow range peak emission of 740 nm and a BP filter of 706/95 nm for bright-field light collection. A Prior NanoScan Z 100 mm piezo z stage (ZEISS) was used for fast acquisition. Optical control images of field of ring (Argolight) and dark current were acquired daily at the start of each data acquisition to monitor microscope performance. Laser power was measured monthly and the corresponding percentage was adjusted to consistently expose the sample to the same laser power. This dataset is available for download at https://open.quiltdata.com/b/allencell/tree/aics/NPM1_single_cell_drug_perturbations/

**Table 1.** List of drugs and concentrations used in the perturbed nucleolar GC dataset.

| Drug | Manufacturer | ID | Concentration(s) |
| --- | --- | --- | --- |
| Actinomycin D | Selleckchem | S8964 | 0.5µg/mL |
| BIX 01294 | Selleckchem | S8006 | 1μM |
| Bafilomycin A | Millipore Sigma | SML1661 | 0.1μM |
| Brefeldin | Selleckchem | S7046 | 5μM |
| Chloroquine | Medchemexpress | HY-17589A | 40μM |
| H89 | Selleckchem | S1582 | 10μM |
| Jasplakinolide | Millipore Sigma | 420127 | 50nM |
| Latrunculin A1 | Millipore Sigma | 428026 | 0.1μM |
| Monensin A | Medchemexpress | HY-N0150 | 1.1μM |
| Nocodazole | Medchemexpress | HY-13520 | 0.1μM |
| Paclitaxel | Selleckchem | S1150 | 5μM |
| Rapamycin | Selleckchem | S1039 | 1μM |
| Roscovitine | Selleckchem | S1153 | 5μM and 10μM |
| Rotenone | Medchemexpress | HY-B1756 | 0.5μM |
| Staurosporine | Selleckchem | S1421 | 1μM |
| Torin-2 | Selleckchem | S2817 | 1μM |

##### 1.4.1 Cell health assessment

We assessed cell health at 4 and 24 hours after drug treatment using for each drug both the AICS-57 (WTC-11 hiPSC endogenously tagged for mEGFP-NMP1) and AICS-61 (WTC-11 hiPSC endogenously tagged for mEGFP-HIST1H2BJ) cell lines. FOVs of this cell line were visually inspected to determine the extent of cell death induced by each drug. If cell death at either 4 or 24 hours was approximately 50% more prevalent than compared to the control, then cells were classified as unhealthy after 2 hours. Otherwise cells were classified as healthy. Results from this assessment are summarized in the last column of the table shown in Figure 7b.

### 2. Input Data Preprocessing for Image Models

#### 2.1 Punctate structures

##### 2.1.1 cellPACK synthetic dataset

Packing results were voxelized into images of size 238x472x472 voxels. The z-coordinate of these images was padded with zeros to be the same size as X and Y, and the resulting images were downsampled to 118x118x118 voxels via block reduce operation with block size of 4x4x4 voxels and then used as input for image-based models.

##### 2.1.2 DNA replication foci dataset

3D raw fluorescence intensity single-cell images of DNA replication foci were masked, centered, and aligned by the corresponding nuclear segmentation dilated by 8x8x8 voxels. Images were cropped and then padded to the largest nuclear bounding box in the dataset. Images were then padded and resized to 118x118x118 voxels. Images were globally contrast adjusted to be within the intensity range 0 to 6000, which was empirically determined to remove dead pixels present in a few images and scaled per image using *monai.transforms.ScaleIntensity*^60^ to be in the range (0,1).

##### 2.1.3 Expanded dataset of punctate structures

Similar preprocessing was applied to a subset of punctate structures from the WTC-11 hiPSC Single-Cell Image Dataset v1^1^, including DNA replication foci, histones, nuclear pores, nuclear speckles, cohesins, peroxisomes, endosomes and centrioles. However, the images of cytoplasmic structures (peroxisomes, endosomes, and centrioles), were masked by the cell membrane segmentation, instead of nuclear segmentation. Images were contrast adjusted using structure-specific intensity ranges reported in^1^. Images were finally scaled per image using *monai.transforms.ScaleIntensity*^60^ to be in the range (0,1). The preprocessing code used to generate this dataset is available at https://github.com/AllenCell/benchmarking_representations/tree/main/br/data/preprocessing/image_preprocessing.

#### 2.2 Polymorphic structures

##### 2.2.1 Nucleoli (GC) dataset

Segmentations of nucleoli (GC) available in the WTC-11 hiPSC Single-Cell Image Dataset v1^1^ were masked by corresponding nuclear segmentations. We used a hole filling algorithm to fill in holes in the segmented images that were then converted into 3D meshes for subsequent preprocessing. Meshes were downscaled to fit within a cube of size 32x32x32 voxels using a global scaling factor to preserve the relative scale of nucleoli in learned representations. For segmentation models the downscaled meshes were voxelized to create binary images. For SDF models the downscaled meshes were used to compute signed distance field images that were clipped to be in the range (-2,2).

##### 2.2.2. Expanded dataset of polymorphic structures

Segmentation of the nucleolar GC, nucleolar DFC, Golgi and lysosomes (available in the WTC-11 hiPSC Single-Cell Image Dataset v1) were masked by either nucleus or cell mask if the structure localizes to nucleus (nucleoli) or cytoplasm (Golgi and lysosomes). Subsequent preprocessing followed Section 2.2.2, except 3D meshes were downscaled on a per cell basis based on the cell’s intracellular structure bounding box. This downscaling avoids losing small nuclear structures given the large bounding box of cytoplasmic structures.

##### 2.2.3 Perturbed nucleoli (GC) dataset

We used the *Allen Cell & Structure Segmenter*^58^ to segment raw fluorescence intensity fields of view (FOVs) of perturbed nucleoli (GC) (section 1.4). Nuclear segmentations for each FOV were produced by applying a UNet model trained on the WTC-11 hiPSC Single-Cell Image Dataset v1^1^ to predict 3D nuclear segmentations from brightfield images. We manually selected nuclear segmentations in each FOV that covered the corresponding the entirety of the nucleoli signal. The selected masks were used to generate single-cell images and they were processed as described in Section 2.2.1.

### 3. Input Data Preprocessing for Point Cloud Models

#### 3.1 Punctate structures

##### 3.1.1 cellPACK synthetic dataset

The list of N=256 centroids of spheres packed by cellPACK was extended to 2,048 points by adding a small jitter to each input point cloud 8 times. This jitter was clipped at a value of 0.2, and the typical range of XYZ coordinates was -10 to 10. This was then used as the 3D point cloud input. To improve reconstruction quality, this augmentation process was repeated 10 times for each input. Details regarding the jitter augmentation are described in section 5.2.

##### 3.1.2 DNA replication foci dataset

We started by applying the same preprocessing used in the DNA replication foci dataset described above for image-based models, except for the last linear scaling step. Instead, we used an exponential function *e*^λ(*skewness*inttensity*)^, where the skewness is a statistic that indicates the deviation of a distribution from a normal distribution. The scaled images were then normalized to obtain a probability density. The coefficient λ is an intracellular-specific scale factor that was empirically determined based on the visualization of sampled points from random images for each intracellular structure. We used λ = 100 for DNA replication foci images. The resulting probability density was used to sample a point cloud with N=20,480 4D (XYZ+intensity) points. Sparse point cloud versions with N=2,048 points were sampled from the original point cloud during training. Point clouds were centered according to the nuclear mask and the intensity coordinate was scaled using by 0.1 to match the magnitude of the spatial coordinates.

#### 3.1.3 Expanded dataset of punctate structures

We started by applying the same preprocessing used in the expanded dataset of punctate structures described above for image-based models, except the last linear scaling step. We again used an exponential function *e*^λ(*skewness*inttensity*)^, with λ = 100 for nuclear structures, and λ = 500 for cytoplasmic punctate structures. The scaled images were then normalized to obtain a probability density. We followed the same procedure described above for DNA replication foci to sample point clouds for each of these punctate structures. The intensity co-ordinate was then normalized using structure specific contrast ranges.

#### 3.2 Polymorphic structures datasets

##### 3.2.1 Nucleoli (GC) dataset

For each single-cell nucleoli (GC) image, we started by sampling a point cloud of N=8,192 points from its corresponding 3D mesh generated as described in Section 2.2.1. A second point cloud with N=20K points is sampled during training from the corresponding SDF volume, yielding a 4D point cloud consisting of SDF value + XYZ coordinates.

##### 3.2.2. Expanded dataset of polymorphic structures

Each single polymorphic structure image underwent a similar process. First, a point cloud of 8,192 points was sampled from the corresponding 3D mesh (see Section 2.2.2). During training, an additional 20K points were sampled from the SDF volume, producing a 4D point cloud (SDF value + XYZ coordinates).

##### 3.2.3 Perturbed nucleoli (GC) dataset

The perturbed nucleoli (GC) dataset followed the same sampling strategy. For each single-cell nucleoli (GC) image, an initial 8,192 point cloud was sampled from the 3D mesh generated as described in section 2.2.3. During training, another point cloud containing 20K points was drawn from the SDF volume, yielding a 4D point cloud (SDF value + XYZ coordinates).

### 4. Model Background

Deep learning for feature extraction has been shown to be powerful in the context of cell biology, in particular, for analyzing images in 2D^61–63^. Despite its success, feature interpretability and generalizability to unseen image data continues to be a major challenge^64,65^. To alleviate some of these problems, it has been shown that imposing additional constraints corresponding to prior biological knowledge to models helps to reduce the space of admissible solutions and improve the likelihood that the learned features can be useful for scientific discovery^66^.

One approach for imposing extra constraints informed by prior knowledge is the integration of known symmetries into the learning process. This has been the guiding principle for the field of geometric deep learning, a subset of machine learning that aims to exploit geometric principles and avoid the curse of dimensionality associated with learning generic functions in high-dimensional spaces^10^. While identifying symmetries in real-life datasets is not always straightforward, enforcing that learned representations are invariant or equivariant under simple Euclidean geometric transformations of the input data has been shown to improve data efficiency and generalization in fields like protein structure prediction ^67^, medical image analysis ^68^, and cell biology ^44^. The distinction between invariant and equivariant representations is of note. While the first type of representation stays the same when the input data is subject to some transformation, the second changes predictably and equivalently. Rotation is a natural choice of geometrical transformation to extract features that do not depend on the object’s orientation.

#### 4.1 Image Models

To implement 3D rotation invariant image autoencoders, we used image encoders equivariant to the group of 3D rotations (*SO3 group*) using R^3^ steerable kernels as described in Weiler et al.^69^ and implemented in the *escnn* library ^70^. Compared to conventional convolutions, R^3^ steerable kernels are equivariant under rotations in R^3^. We used scalar fields to learn invariant scalar features in R^3^, and vector fields to learn equivariant vector features in R^3^. We used vector features to reconstruct the 3D rotation matrix as described in Deng et al.^14^ and Winter et al.^77^.

We used seven layers of steerable kernels with an equal number of hidden scalar fields using trivial representations and vector fields using irreducible representations. Using a (filter, stride, kernel size) convention, the convolutions were (8, 1, 3), (16, 1, 3), (32, 2, 3), (64, 2, 3), (128, 2, 3), (512, 2, 3), (N, 1, 1), where N was the size of the latent dimension. In the final layer we used N scalar fields and 2 vector fields. Each convolutional block also included a batchnorm and ReLU activation^71^. We used average pooling in the last 5 layers and checked that this did not break equivariance (Fig. S2). We spatially pooled the scalar embedding in the final layer to get the final N dimensional rotation invariant latent embedding. We used a bottleneck size of 512 for polymorphic structures and 256 for punctate structures.

The decoding function was a conventional CNN decoder with 6 layers of convolutions. We used upsampling blocks with a scale factor of 2 in between convolutions. Using a (filter, stride, kernel size) convention, the convolutions were (512, 1, 3), (256, 1, 3), (128, 1, 3), (64, 1, 3), (32, 1, 3), (16, 1, 3). We rotated the canonical reconstruction with the rotation matrix computed from the vector representation. We used a cylinder mask using *escnn.nn.modules.masking_module.build_mask* to mask reconstructions and reduce interpolation artifacts. We set the background value to 0 for segmentations, and 2 for clipped SDF images where the maximum value was 2 and positive values were located outside the object. We used the same settings with classical autoencoders by swapping out equivariant convolutions with regular convolutions and keeping other details the same.

##### 4.1.1 Masked autoencoders using vision transformers

We also trained masked autoencoders (MAE) using vision transformers^72^ in two stages as an alternative to the *vanilla* autoencoders described above. We performed this training in two stages. First, we pretrained a MAE^73^ using a ZYX patch size of (2,2,2), a mask ratio of 0.75, and learnable positional embeddings. The encoder was made up of 8 identical transformer blocks, each with 4 heads and an embedding dimension of 256. The decoder had 2 layers with 8 heads and an embedding dimension of 192. We then employed a second phase of training with a mask ratio of 0 (i.e. all image patches are visible to the encoder) where we froze the MAE-trained encoder and trained a freshly initialized decoder to reconstruct the input image. We trained all models with a mean squared error (MSE) loss.

#### 4.2 Point cloud models

To implement 3D rotation invariant point cloud autoencoders, we used a 3D rotation equivariant point cloud encoder using vector neurons (VN^11^), which lifts classical neurons to 3D vectors resulting in 3D vector representations. VN layers are equivariant to rotations by construction, and have been shown to outperform other equivariant architectures for tasks like classification, segmentation, and reconstruction. We incorporate VN layers into a Dynamic Graph CNN (DGCNN)^74^ backbone for point cloud encoding. DGCNN uses network modules called EdgeConvs to perform CNN-like local neighborhood feature extraction. These EdgeConvs can be stacked to extract global features^74^. Dynamic graphs are computed by constructing k-nearest neighbor graphs on points. We used k=20 based on previous works as a balance between computational complexity and local structure information^45^. We concatenated the cross-product of the neighbor features and input points as well as the input points themselves to the hidden representation. As described in section 3, we included raw image intensity in addition to XYZ coordinates in some cases to generate 4D point clouds. This coordinate was included with the same vector orientation as the XYZ coordinates, and thus remains equivariant under rotations in R^3^. For the cellPACK dataset, we used a 3D point cloud as input. We used 6 convolutional blocks where each block comprises a VN Linear layer and a VN Leaky ReLU layer. We collated intermediate outputs before a final 1D convolution. We took the norm of the final vector embedding to get a rotation invariant representation. We also trained classical point cloud autoencoders with DGCNN encoders as described in Vries et al. ^45^, where VNLinear and VN LeakyReLU layers are replaced with edge convolutions and ReLU layers.

##### 4.2.1 Decoder for punctate structures

We reconstructed the rotation invariant representation for punctate structures using a folding net decoder^75^. This decoder concatenates the latent embedding with source points sampled from a template shape, and then applies two folding operations with ReLU activations interleaved in between to reconstruct a point cloud. We used a 2D plane as a template in all cases except for the cellPACK synthetic dataset, where a sphere was used as a template. Next, we used the learned rotation matrix from the vector embedding to re-orient the canonical reconstruction. We optimized the model using an earth mover’s distance^76^.

##### 4.2.2 Decoder for polymorphic structures

We reconstructed the rotation equivariant representation computed using vector neurons via an inner product decoder^11^ to get signed distance function values at query points. These query points correspond to the points of the second point cloud sampled from the SDF volume as described in section 3.2. The model is optimized using an L1 loss on the SDF values. We took the norm of the embeddings after training to compute a rotation invariant representation. This relaxes the generative nature of the model as we are no longer able to obtain a rotation invariant reconstruction. This trades off the ability to generate rotation-invariant predictions with reconstruction quality, deemed necessary for the more complex shapes of the polymorphic structures. Instead, we visualize the closest real examples to different map points in representation space (Fig. 5, 6).

### 5. Model Training

#### 5.1. Train, validation and test splits

We used a train/valid/test split ratio of 70%, 15%, 15% respectively across all datasets. For the expanded polymorphic dataset, we stratified the split using the structure class. For the perturbed nucleoli (GC) dataset, we stratified the split by drug.

#### 5.2 Jitter augmentation

We trained the rotation invariant point cloud models with an augmented version of the DNA replication foci dataset (N=2,420) and the synthetic dataset from cellPACK (N=254) because of the small size of these datasets. This augmentation was done by adding jitter to each input point cloud (with 2,048 points) 10 times during training. The jitter was added via Gaussian noise with a standard deviation of 0.1. The noise was clipped at a value of 0.2. The typical range of XYZ coordinates was -10 to 10. Adding augmentations helped improve the quality of reconstructions (Fig. S4).

### 6. Multi-metric Model Benchmarking

We computed three sets of metrics to quantitatively benchmark our trained models (Fig. S1a-b). Efficiency metrics including model size, inference time, and emissions to assess ease of inference and training. Generative metrics included reconstruction error and evolution energy^7^ to assess the quality of generated shapes. Finally, representation expressivity metrics evaluated classification/regression scores for biologically-relevant features, rotation invariance error, and interpolation distances in embedding space.

#### Efficiency

*Model size* is the number of parameters in the model, *inference time* is the time to run GPU inference on a single input, and *carbon emissions* is an estimate of hardware electricity power consumption in kWh. All measurements use a single A100 GPU.

#### Generative capacity

*Reconstruction error* is the average test set reconstruction error using the Jaccard similarity score for images and Chamfer distance for point clouds, respectively. More details on classification and regression calculations can be found in section 6.1. *Evolution energy* is the normalized energy of deformation from one shape to another averaged across many random test set pairs in a dataset (Fig. S1b). Here, the normalized energy of deformation is computed as the sum of the reconstruction error between an interpolated shape reconstruction and initial and final reconstructions, divided by the reconstruction error between initial and final reconstructions. This energy is computed across 10 interpolations between two given shapes.

#### Expressivity

*Rotation invariance error* is computed as the norm of the Euclidean distance between the embedding of a test set shape and its rotated version, divided by the sum of the norms of the two embeddings. The metric aims to quantify how much the embeddings change in representation space with rotation, while accounting for the size of the embedding space itself via

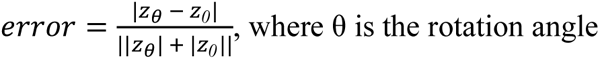

By dividing by the sum of the norms of the two embeddings, we establish their size relative to the origin. We computed this metric for four 90 degree rotations of the input in the XY plane. This is then averaged across many test set examples. *Average interpolation distance* is computed as the distance to the closest real example in representation space for interpolations between random test set examples (Fig. S). We did this for 10 interpolations between randomly sampled pairs of test set examples and report the average across pairs and interpolations. *Compactness* is an intrinsic dimensionality measure calculated using the Levina-Bickel score^19^. All metrics were z-scored across models for polar plot visualization. The sign of all metrics except for classification and regression scores were flipped to ensure that a higher value indicates better performance. *Classification* and *regression scores* are the respective cross-validated test set accuracies and R^2^ values for selected features. More details on classification and regression calculations can be found in section 6.2.

#### 6.1 Reconstruction error

##### Punctate structures

We used the Chamfer loss to compute the reconstruction error of image- and point cloud-based models. Because this loss is normally applied to point clouds, we convert inputs and respective reconstructions from image models into point clouds by sampling points using the exponential sampling function described in section 3.1.

##### Polymorphic structures

We used the Jaccard similarity score on binary masks to compute the reconstruction error of image-(segmentation and SDF) and point cloud-based models. For segmentation models, we used Otsu thresholding to binarize each reconstruction and extracted a mesh using marching cubes. Next, we upscaled the mesh and voxelized it into a 3D binary mask. To generate a binary mask for SDF images used as input for SDF models and their reconstructions, we first used the marching cubes algorithm to extract a zero-level set mesh. Then we upscaled the mesh and voxelized it to create a 3D binary mask. For SDF point cloud models, we used vertices of a 32x32x32 grid as XYZ query points to perform inference along with ground truth surface point clouds to obtain SDF reconstruction images. We then used the marching cubes algorithm to extract a zero-level set mesh from each SDF image, upscaled the mesh, and voxelized it into a 3D binary mask.

#### 6.2 Classification and Regression Scores

We normalized embeddings using *StandardScaler* from scikit-learn^77^. For classification accuracies, we used a logistic regression classifier with class weights and reported test set classification accuracies with stratified k-fold cross-validation with five splits. For regression scores, we used a linear regression model and reported test set R^2^ with repeated k-fold cross validation with five splits and 20 repeats.

Classification and regression targets were designed appropriately. For the cellPACK synthetic dataset, we reported cross-validated accuracy for classifying the six packing rules. For the DNA replication foci dataset, we reported cross-validated top 2 classification accuracy for 8 interphase cell cycle stages and cross-validated accuracy for manually annotated outlier labels including dead cells and cells with no fluorescence. For punctate structures from the WTC-11 hiPSC Single-Cell Image Dataset v1, we reported cross-validated structure classification accuracy for 7 structures and cross-validated classification accuracy for cell cycle stages including interphase, prophase, early prometaphase, prometaphase/metaphase and anaphase/telophase. For the nucleoli (GC) dataset, we reported cross-validated classification accuracy of thresholded number of pieces of nucleoli present in the segmentation, including 1, 2, 3, 4 and >=5 piece nucleoli. For this dataset, we also reported an average test set R^2^ for regression of mean and standard deviation of piece volume and area, and an average test set R^2^ for regression of mean and standard deviation of centroid-centroid distances between pieces. For the polymorphic structure dataset including nucleoli (GC and DFC), lysosomes and Golgi, we reported cross validated classification accuracy for structures, and regression scores for volume and area of pieces, and distance between pieces similar to the nucleoli (GC) dataset.

### 7. Data Analysis

#### 7.1 Principal component analysis (PCA)

We fit PCA to the learned rotation invariant representations using *sklearn.decomposition.PCA* with “auto” SVD solver and the number of components equal to the size of the embedding space. We then performed inverse PCA on different map points (-2σ, 0, 2σ) for different principal components (PCs). We used the inverse PCA reconstruction as input to the decoder of the representation learning model to visualize the reconstructed image/point cloud.

#### 7.2 Archetype analysis

Archetype analysis approximates the convex hull of data points by identifying extremal points^20^. We used an implementation of the Frank-Wolfe algorithm for archetype analysis^78^. The number of archetypes for each dataset was chosen based on an expected number of clusters in each application.

#### 7.3 Nuclear volume binning for nucleoli (GC) and DNA replication foci dataset

We clipped the nuclear volume distribution to be within the 2.5% and 97.5% range for both DNA replication foci and nucleolar GC datasets. Next, we binned the data into equal sized bins of 121 µm^3^. The bins were <390 µm^3^, 390-533 µm^3^, 533-676 µm^3^, 676-818 µm^3^, and >818 µm^3^.

### 8. Data Availability

The WTC-11 hiPSC single cell image dataset v1 analyzed in this study is available online at https://open.quiltdata.com/b/allencell/packages/aics/hipsc_single_cell_image_dataset. The DNA replication foci dataset analyzed in this study is available online at https://open.quiltdata.com/b/allencell/packages/aics/nuclear_project_dataset_4. The WTC-11 hiPSC nucleoli (NPM1) perturbation single cell image dataset analyzed in this study is available online at https://open.quiltdata.com/b/allencell/tree/aics/NPM1_single_cell_drug_perturbations/. The synthetic dataset of punctate structures generated using cellPACK and analyzed in this study is available online at https://open.quiltdata.com/b/allencell/tree/aics/morphology_appropriate_representation_learning/cellPACK_single_cell_punctate_structure/. The landing page of the GitHub repository associated with this manuscript (https://github.com/AllenCell/benchmarking_representations) has additional information for accessing and processing these datasets.

### 8. Code Availability

Code for all our representation learning models is available at https://github.com/AllenCellModeling/cyto-dl/blob/br_release/.

Config files associated with our models, training scripts, and code for multi-metric benchmarking are available at https://github.com/AllenCell/benchmarking_representations. Our code was all developed in Python. A full list of dependencies is available at https://github.com/AllenCellModeling/cyto-dl/tree/br_release/requirements and https://github.com/AllenCell/benchmarking_representations/blob/main/pyproject.toml. We have released the code with the Allen Institute Software License.

## Supplemental figures

**Figure S1.**
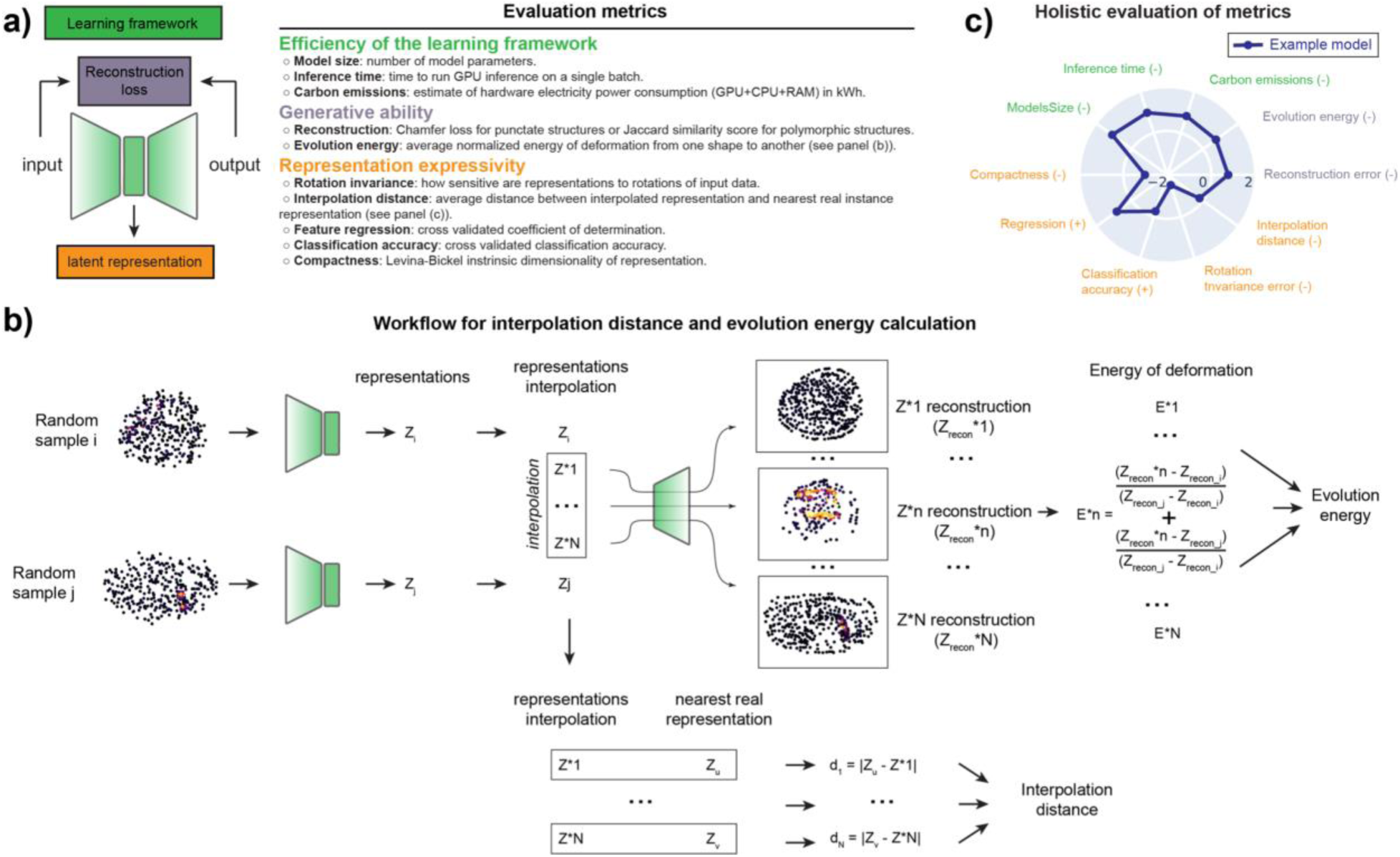
Evaluation metrics for representation learning models. **a)** Overview of different evaluation metrics for quantifying the utility of each representation learning framework. Efficiency metrics include model size, inference time, and carbon emissions. Generative ability metrics include reconstruction error and evolution energy. Representation expressivity metrics include rotation invariance error, interpolation distance, feature regression, classification accuracy, and compactness. **b)** Workflow for interpolation distance and evolution energy calculation. Two samples are drawn from the population randomly, and a linear interpolation is performed on the representations of the two samples. The euclidean distance between an interpolation and the nearest real representation is the interpolation distance. The interpolation distance is averaged across many interpolations to compute the average interpolation distance. Each interpolation is reconstructed using the decoder to obtain a reconstruction. The sum of the reconstruction error between the interpolated reconstruction and the reconstructions of the initial and final shapes normalized by the reconstruction error between the initial and final shape is the energy of deformation^7^. The energy of deformation is averaged across many interpolations to compute the evolution energy. Both evolution energy and average interpolation distance are averaged across many random pairs of samples from the test set. **c)** Holistic evaluation of metrics. Metrics are z-scored across models per metric. Z-scored metrics are visualized using a polar plot by flipping the sign for metrics where lower is better (indicated by a negative sign).

**Figure S2.**
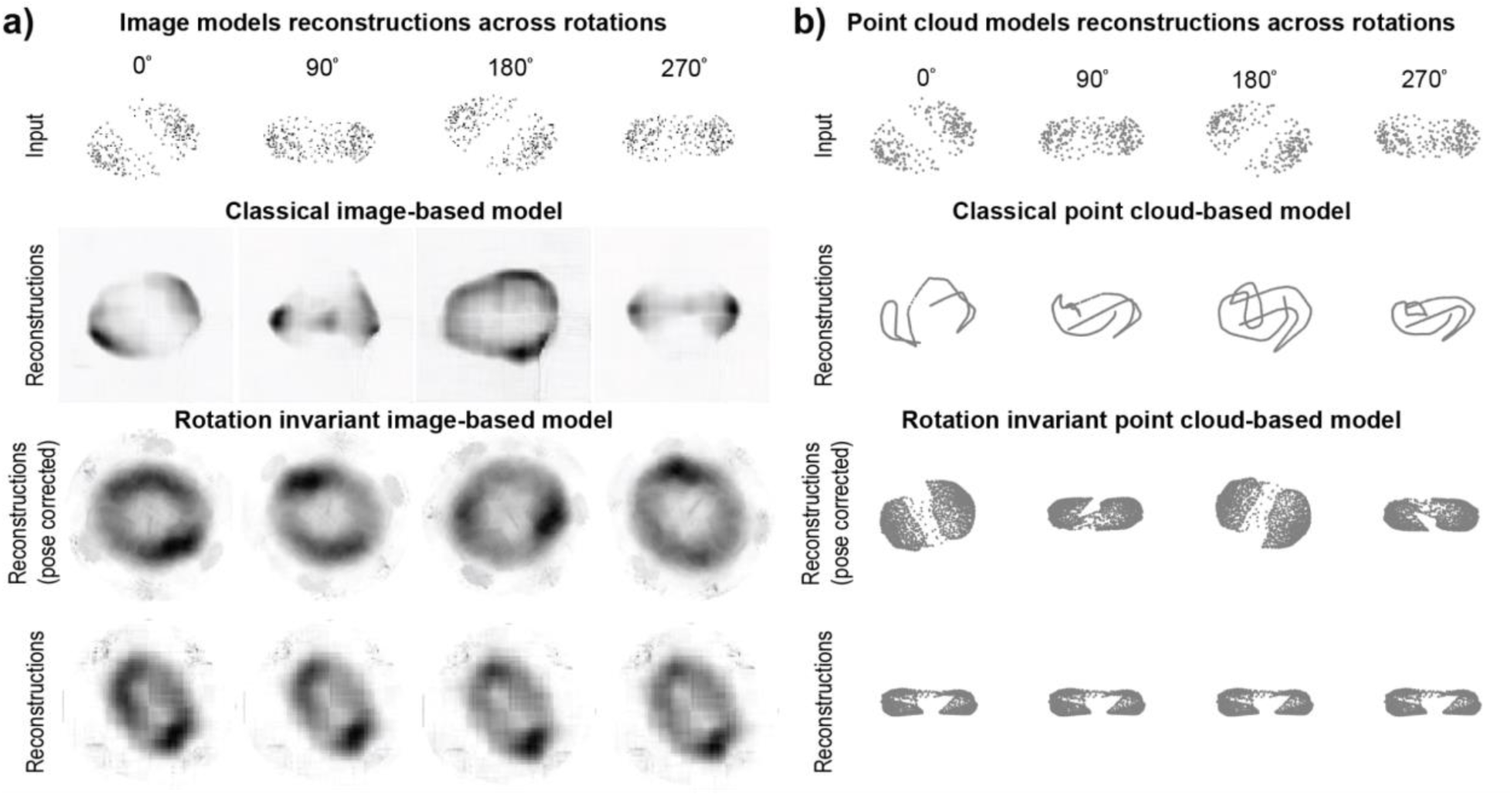
Testing orientation invariance for image and point cloud models for the cellPACK synthetic dataset. **a)** (Top row) Example image input for the planar 45 rule is rotated by four 90 degree rotations. (Middle row) Reconstructions using the classical image model (upper) and rotation invariant image model (lower) for each rotated input. The reconstructions using the rotation invariant model are pose-corrected using the learned rotation angles. (Bottom row) Rotation invariant reconstructions using the rotation invariant image model for each rotated input. **b)** (Top row) Example point cloud input for the planar 45 rule is rotated by four 90 degree rotations. (Middle row) Reconstructions using the classical point cloud model (upper) and rotation invariant point cloud model (lower) for each rotated input. The reconstructions using the rotation invariant model are pose-corrected using the learned rotation angles. (Bottom row) Rotation invariant reconstructions using the rotation invariant point cloud model for each rotated input. All reconstructions shown are max projections in Z.

**Figure S3.**
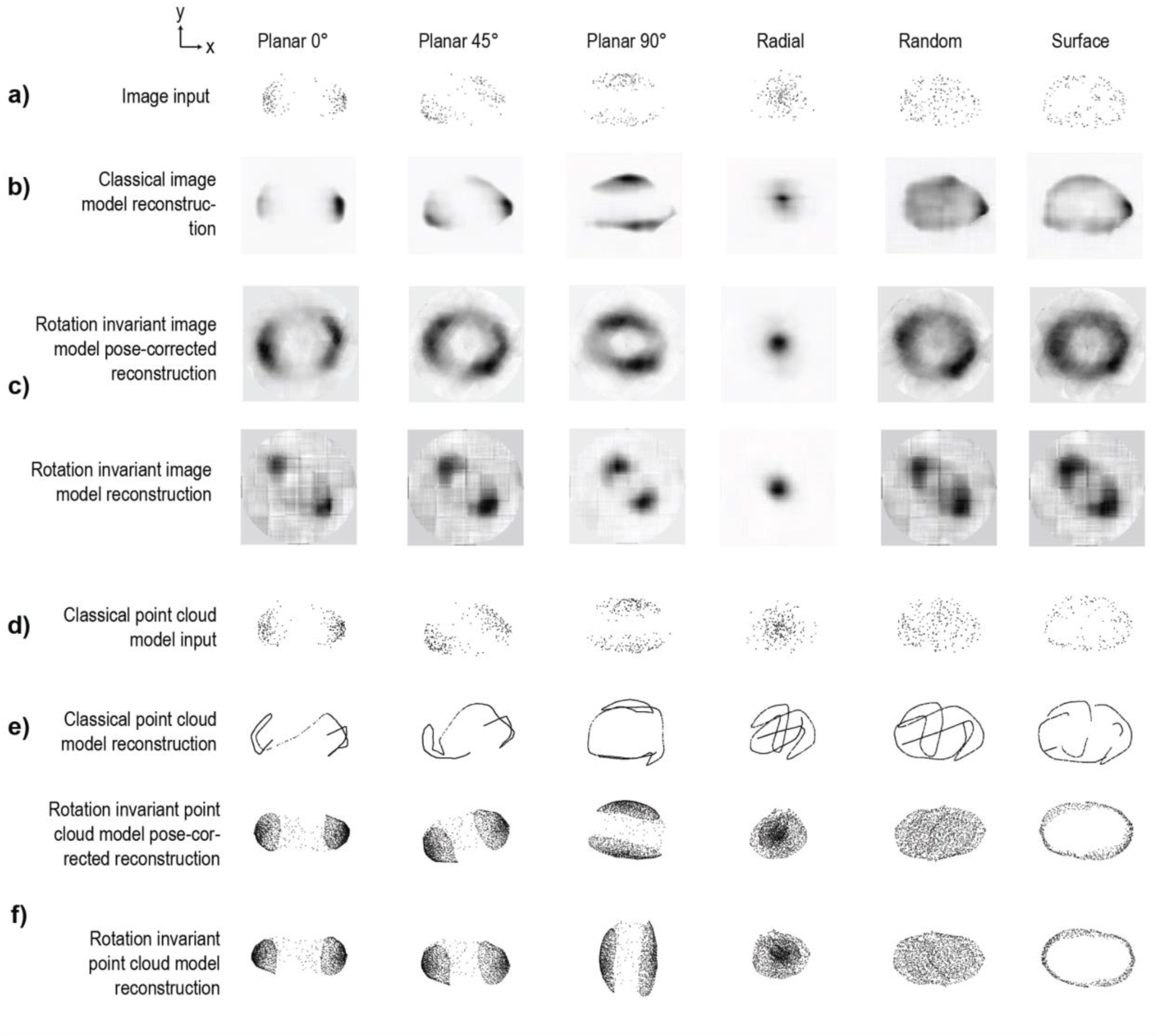
Evaluation of test set model reconstructions for synthetic punctate structures generated using cellPACK. Test set center slice inputs (**a, d)** and reconstructions using **b)** classical image model, **c)** rotation invariant image model, **e)** classical point cloud model, and **f)** rotation invariant point cloud model for each of the 6 packing rules. Both pose-corrected and rotation invariant reconstructions are shown for the rotation invariant models.

**Figure S4.**
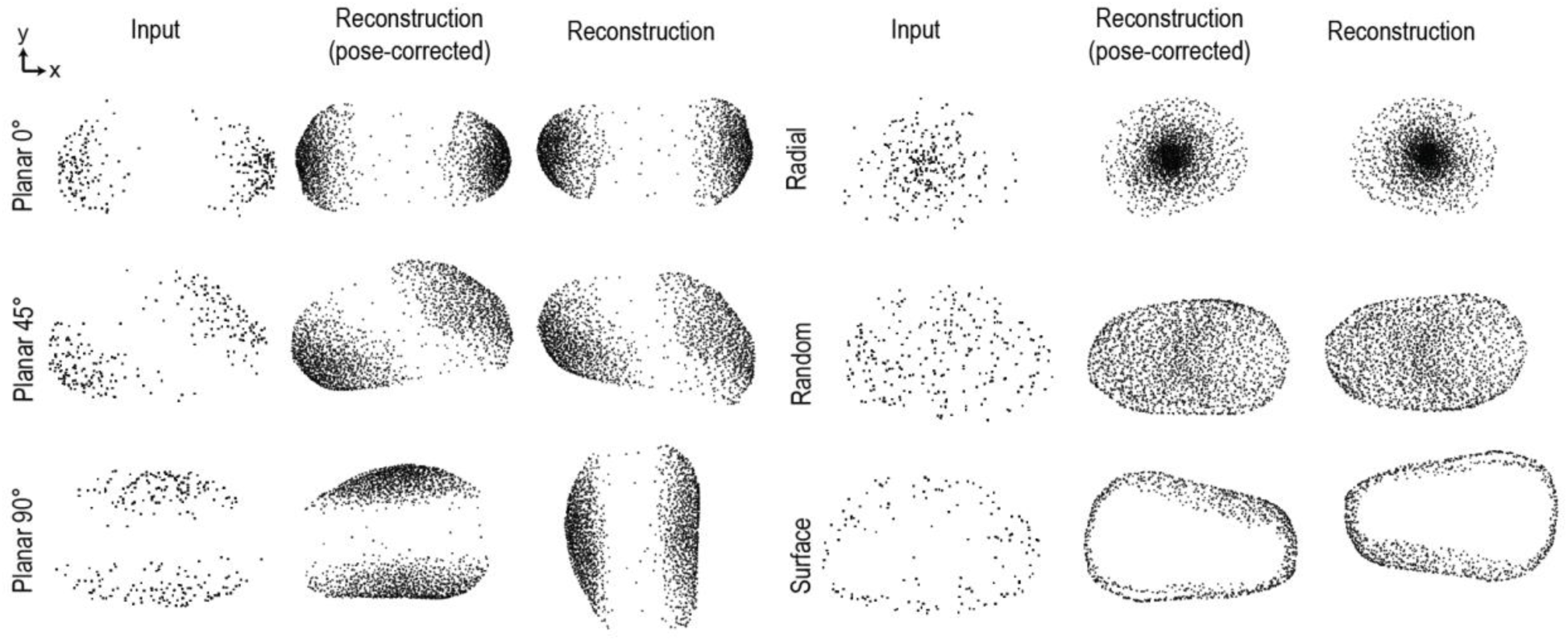
Jitter augmentations slightly improve point cloud model reconstructions for synthetic punctate structures generated using cellPACK. Test set center slice reconstructions using rotation invariant point cloud model with jitter augmentations for each of the 6 packing rules.

**Figure S5.**
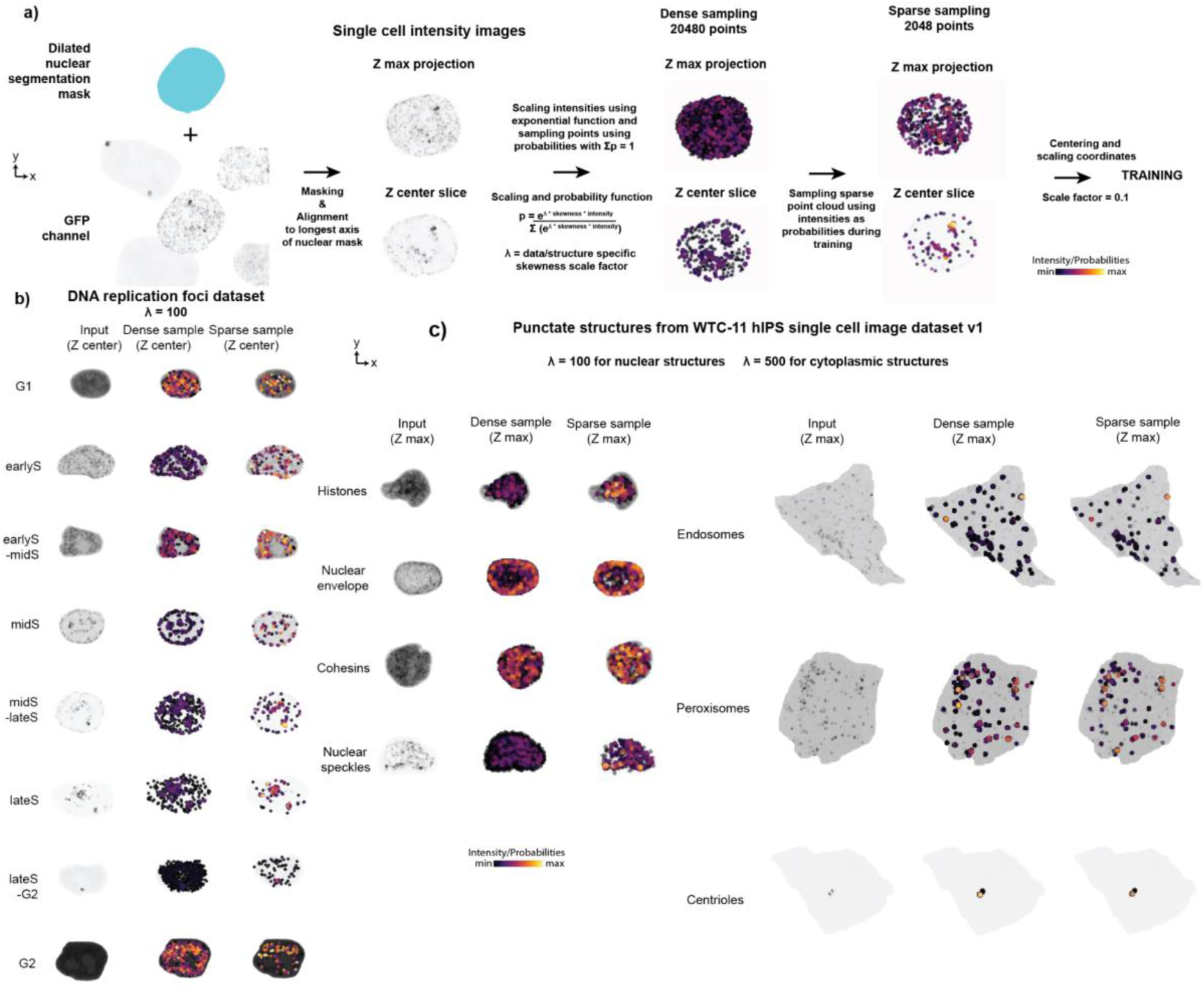
3D image preprocessing into application appropriate inputs for punctate structures. Workflow for generating 4D point clouds from 3D intensity images. **a)** single-cell intensity images are obtained by masking via a dilated nuclear mask (for nuclear structures), followed by alignment to the longest axis of the nuclear mask. Intensities were then scaled using an exponential function and then converted to probabilities. These probabilities were then used to sample a dense 4D point cloud with 20480 points and XYZ + intensity coordinates. During training, a sparse point cloud with 2048 points was sampled from this dense point cloud using the intensities as probabilities. The intensity coordinate was scaled using a scale factor of 0.1 to ensure that intensity values were in the same range as XYZ coordinate values. **b)** Examples of dense sample and sparse sample for each cell cycle stage for PCNA dataset. Shown are center-slice of raw intensity image, center-slice of raw intensity image overlaid with dense sample, and center-slice of raw intensity image overlaid with sparse training sample. **c)** Examples of dense sample and sparse sample for each punctate structure from the WTC-11 hiPSC Single-Cell Image Dataset v1. Structures include histones, nuclear envelope, cohesins, nuclear speckles, endosomes, peroxisomes, and centrioles.

**Figure S6.**
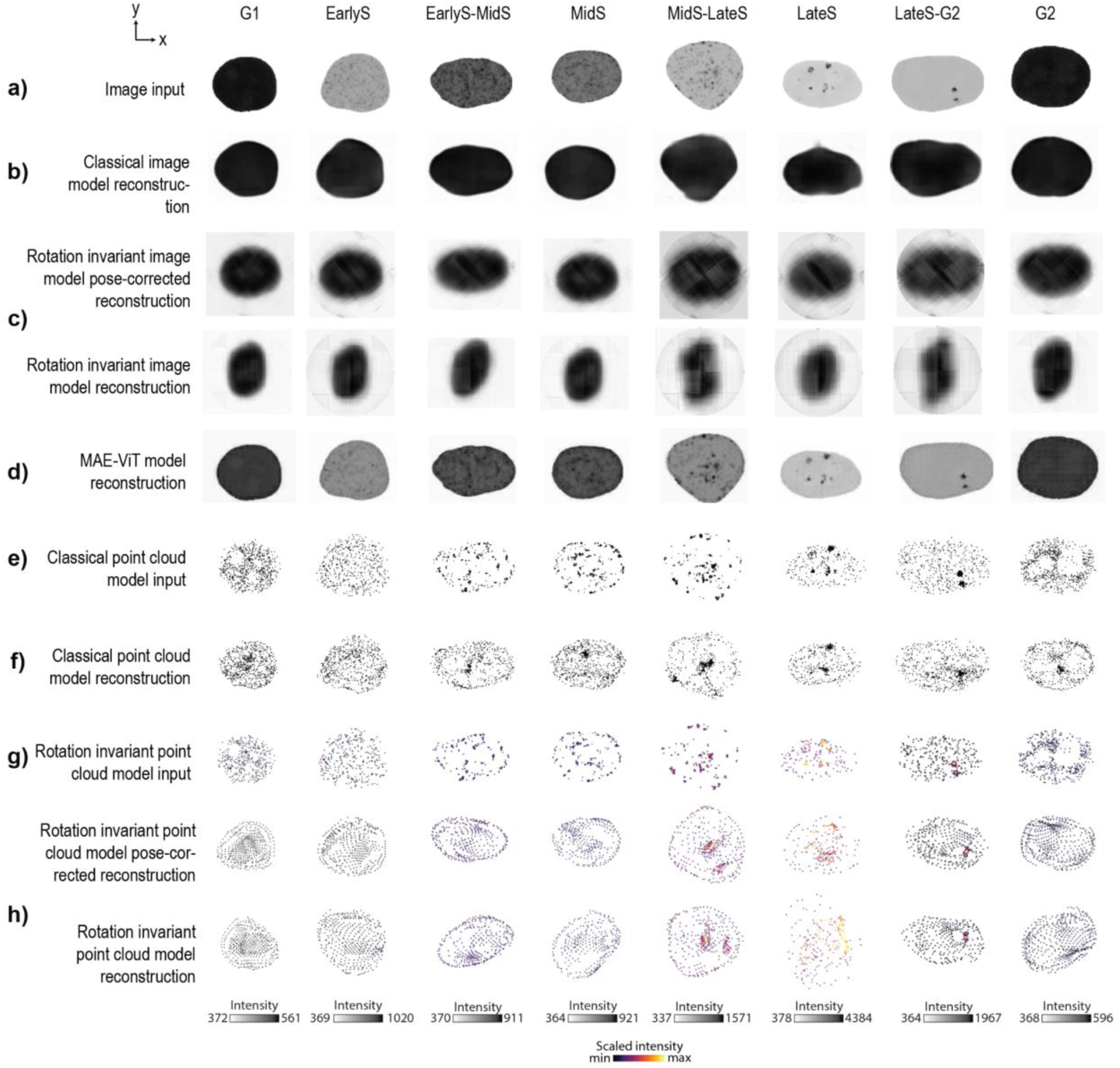
Evaluation of test set model reconstructions for the DNA replication foci dataset. Test set center slice inputs (**a, e, g**) and reconstructions using **b)** classical image model, **c)** rotation invariant image model, **d)** an alternative classical image model via a masked autoencoder with a vision transformer as an encoder (MAE-ViT), **f)** classical point cloud model, and **h)** rotation invariant point cloud model for samples from each of the 8 cell cycle stages. Both pose-corrected and rotation invariant reconstructions are shown for the rotation invariant models.

**Figure S7.**
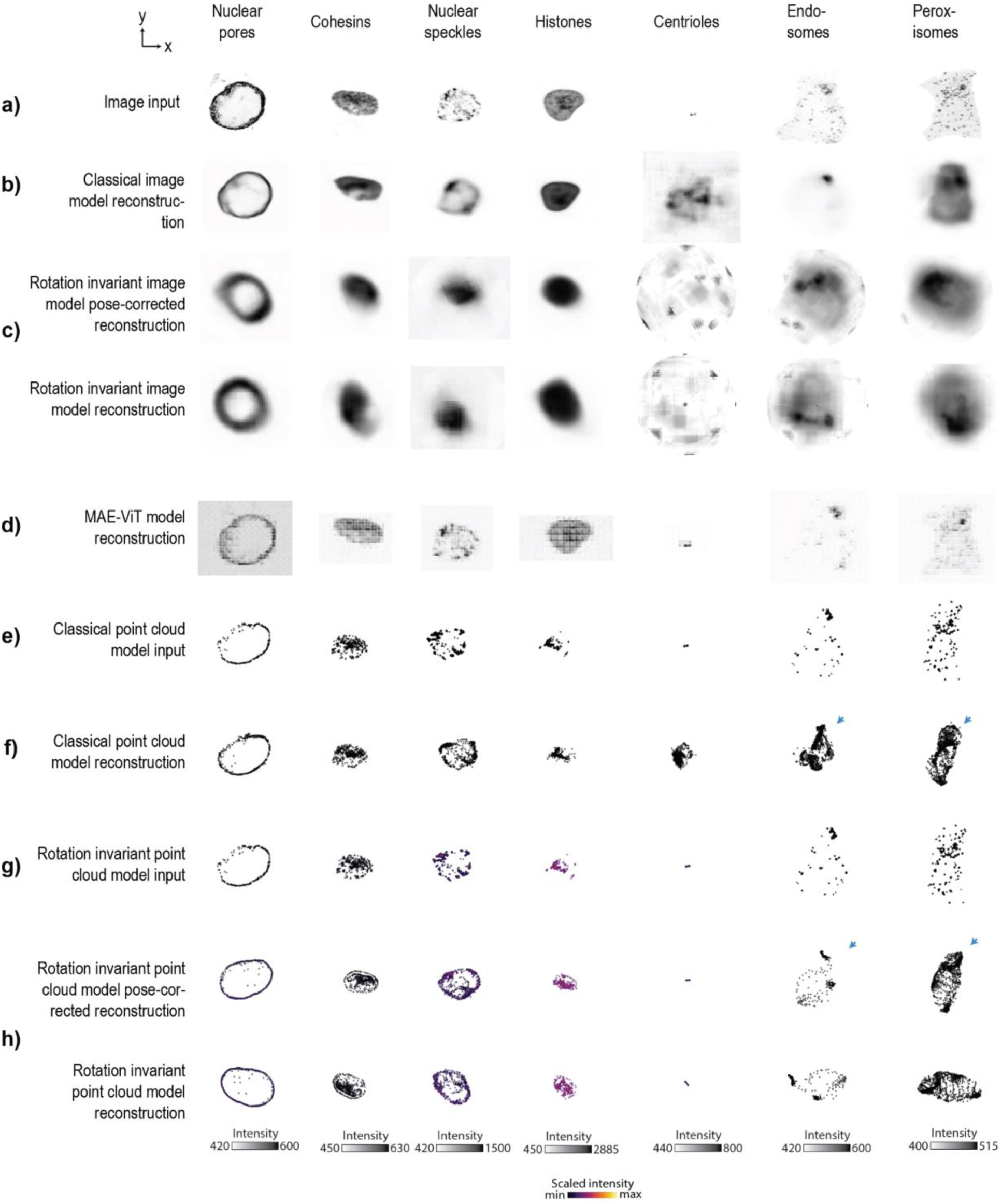
Evaluation of test set model reconstructions for punctate structures from the WTC-11 hiPSC Single-Cell Image Dataset v1. Visualization of test set reconstructions for sampled histones (CellId 721646), nuclear pores (CellId 873680), cohesins (CellId 994027), nuclear speckles (CellId 490385), centrioles (CellId 451974), endosomes (CellId 811336), peroxisomes (CellId 835431). Shown are test set inputs (**a, e, g**) and reconstructions using **b)** classical image model, **c)** rotation invariant image model, **d)** an alternative classical image model via a masked autoencoder with a vision transformer as an encoder (MAE-ViT), **f)** classical point cloud model, and **h)** rotation invariant point cloud model for each structure. Both pose-corrected and rotation invariant reconstructions are shown for the rotation invariant models. Reconstructions for nuclear pores, cohesins, and histones are center slices, whereas reconstructions for nuclear speckles, centrioles, endosomes, and peroxisomes are max projections. Spatial distribution artifacts in reconstructions for endosomes and peroxisomes are highlighted with blue arrows.

**Figure S8.**
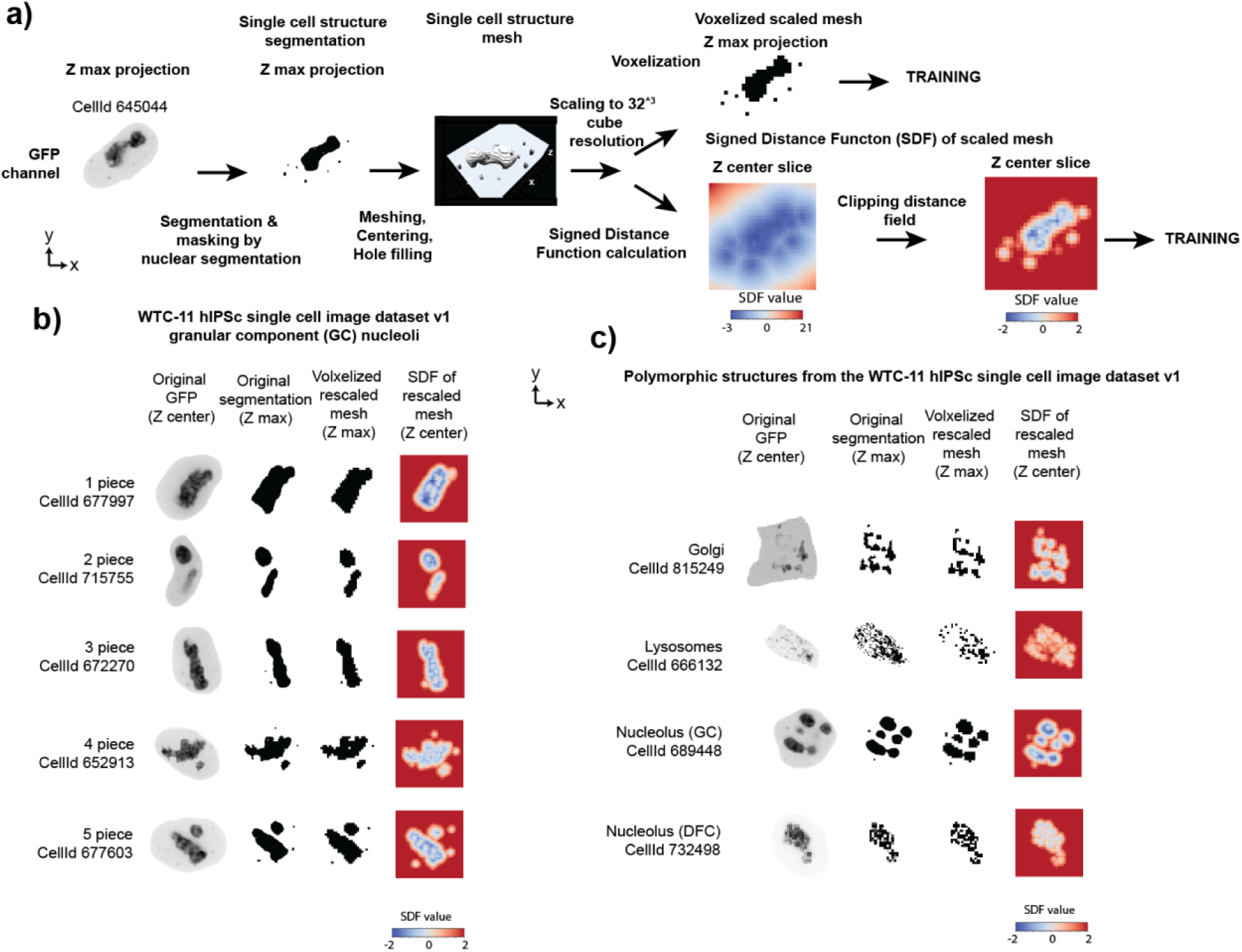
3D image preprocessing into application appropriate inputs for polymorphic structures. **a)** Workflow for computing signed distance function (SDF) images from segmentations. single-cell structure segmentations are masked by nuclear segmentation (for nuclear structures), followed by meshing, centering, and hole filling. The mesh is then rescaled to 32*^3^ cube resolution and then processed to get a signed distance function. Alternatively, the rescaled mesh is voxelized to get a segmentation. SDF is clipped to (-2, 2) range for training image models to focus models on the zero level set. Example shown is for nucleoli (GC). **b)**. Visualization of rescaled segmentation and SDF for examples with different numbers of pieces of granular component (GC) of nucleoli. Shown are center-slices of raw intensity images, max projection of the structure segmentation, max projection of the voxelized rescaled segmentation, and center slice of the rescaled mesh SDF. **c)** Visualization of rescaled segmentation and SDF for other polymorphic structures from the WTC-11 hiPSC Single-Cell Image Dataset v1 including lysosomes, Golgi, GC nucleoli, and dense fibrillar component (DFC) nucleoli.

**Figure S9.**
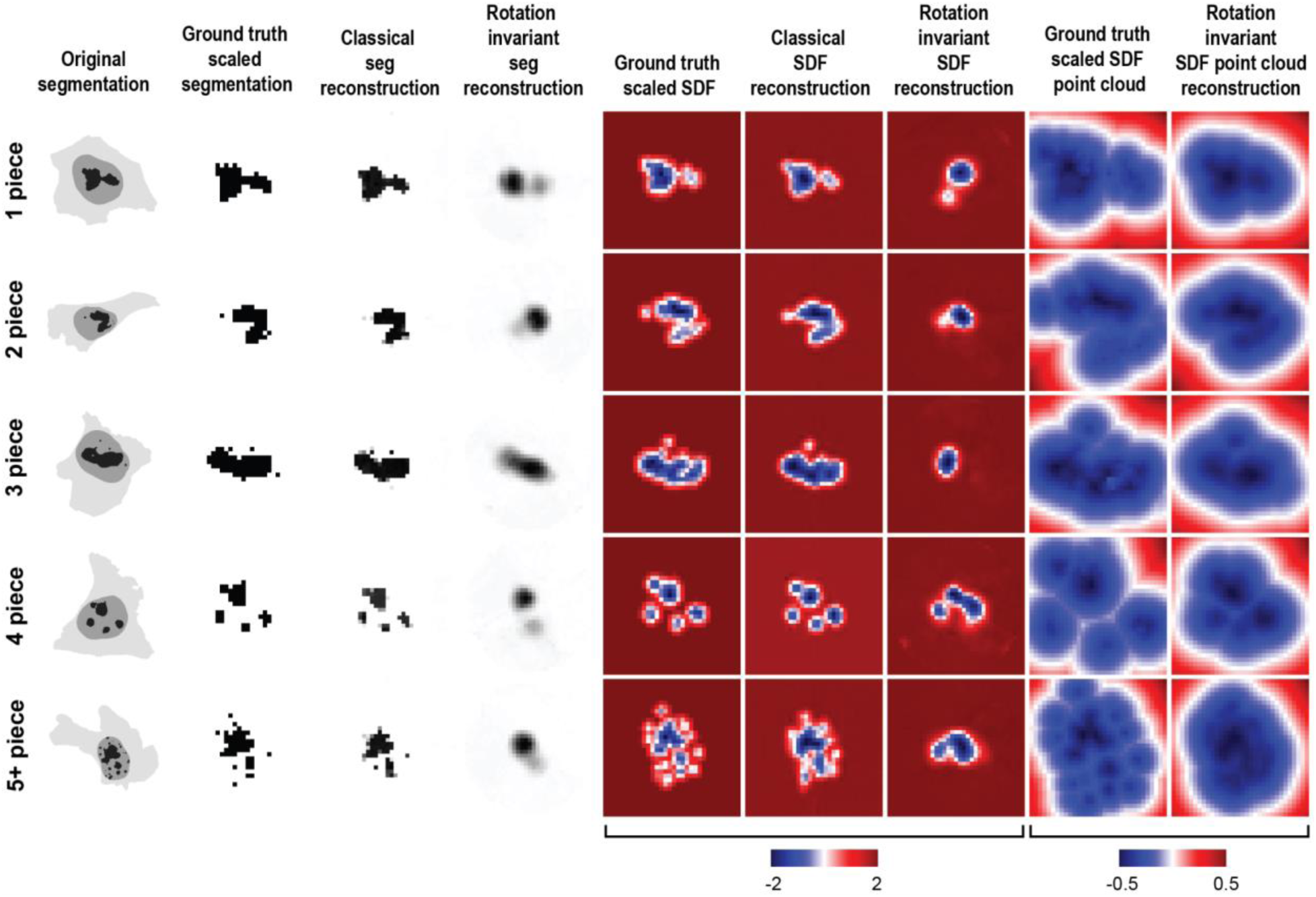
Examples of model inputs and outputs for nucleolar GC dataset. Test set reconstructions across all models for sampled 1 piece (CellId 964798), 2 pieces (CellId 661110), 3 pieces (CellId 644401), 4 pieces (CellId 967887) and 5+ pieces (CellId 703621) examples. Max projections of original structure segmentations overlaid with nuclear and membrane segmentations are shown. Max projections are shown for segmentations, whereas middle slices are shown for SDFs.

**Figure S10.**
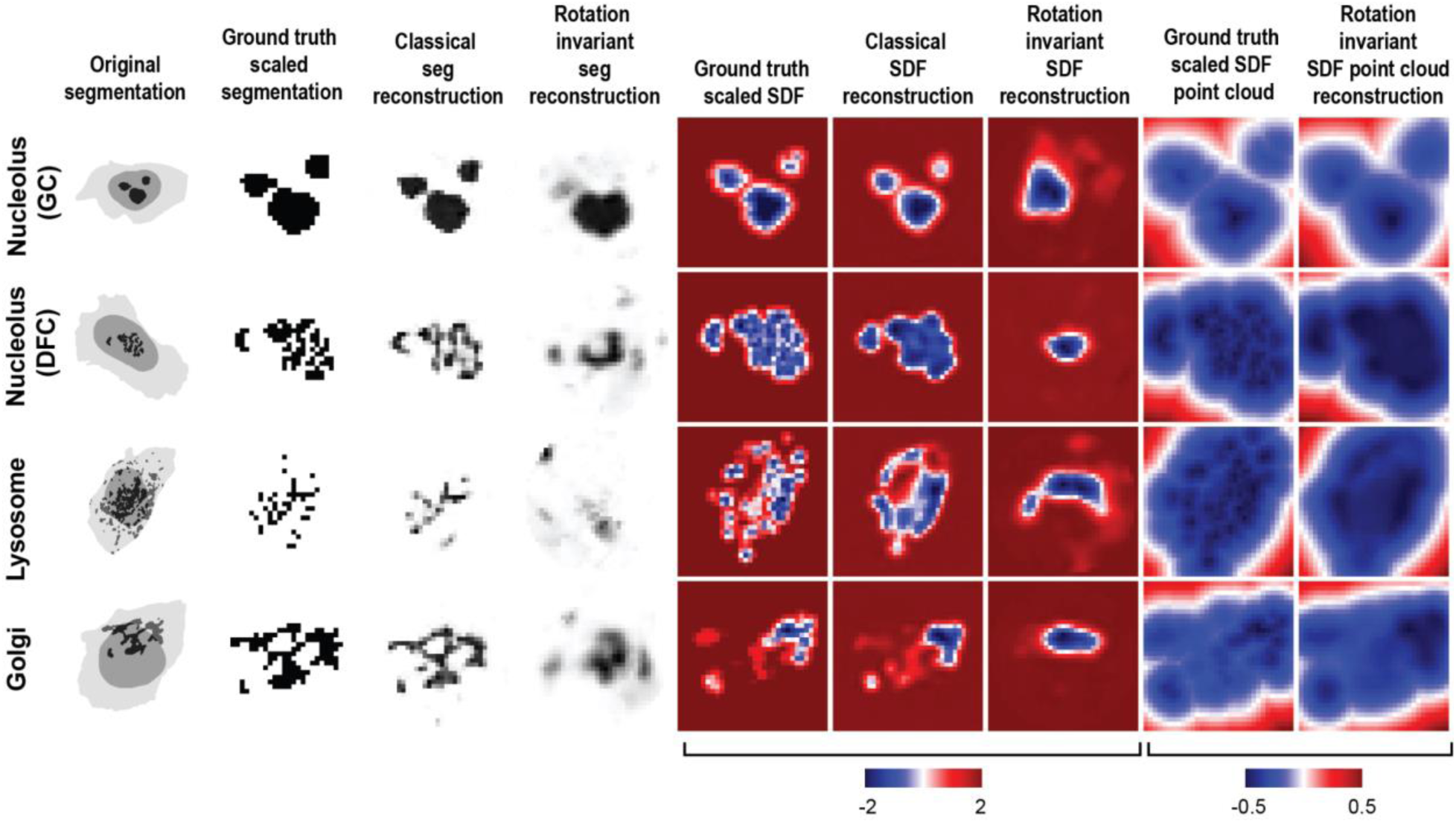
Evaluation of model reconstructions for polymorphic structures from the WTC-11 hiPSC Single-Cell Image Dataset v1. A) Test set reconstructions across all models for sampled nucleoli (GC) (CellId 691110), nucleoli (DFC) (CellId 723687), lysosome (CellId 816468), and Golgi (CellId 800894) examples. Max projections of original structure segmentations overlaid with nuclear and membrane segmentations are shown. Max projections are shown for segmentations, whereas middle slices are shown for SDFs.

## Notes

https://github.com/AllenCell/benchmarking_representations

https://open.quiltdata.com/b/allencell/tree/aics/NPM1_single_cell_drug_perturbations/

https://open.quiltdata.com/b/allencell/tree/aics/morphology_appropriate_representation_learning/cellPACK_single_cell_punctate_structure/

https://open.quiltdata.com/b/allencell/packages/aics/nuclear_project_dataset_4

https://open.quiltdata.com/b/allencell/packages/aics/hipsc_single_cell_image_dataset

### Competing Interest Statement

The authors have declared no competing interest.

### Summary of Updates

We found that some hyperlinks in the text were broken. We have fixed these links in this new version.

